# Mass Spectrometry-based Profiling of Single-cell Histone Post-translational Modifications to Dissect Chromatin Heterogeneity

**DOI:** 10.1101/2024.07.05.602213

**Authors:** Ronald Cutler, Laura Corveleyn, Claudia Ctortecka, Joshua Cantlon, Sebastian Alvaro Jacome Vaca, Dieter Deforce, Jan Vijg, Maarten Dhaenens, Malvina Papanastasiou, Steven A. Carr, Simone Sidoli

**Author notes:** Authors contributed equally to this work (Ronald Cutler, Laura Corveleyn, Claudia Ctortecka). Correspondence to R.C., S.A.C., M.P. and S.S.

## Abstract

Single-cell proteomics confidently quantifies cellular heterogeneity, yet precise quantification of post-translational modifications, such as those deposited on histone proteins, has remained elusive. Here, we developed a robust mass spectrometry-based method for the unbiased analysis of single-cell histone post-translational modifications (schPTM). schPTM identifies both single and combinatorial histone post-translational modifications (68 peptidoforms in total), which includes nearly all frequently studied histone post-translational modifications with comparable reproducibility to traditional bulk experiments. As a proof of concept, we treated cells with sodium butyrate, a histone deacetylase inhibitor, and demonstrated that our method can i) distinguish between treated and non-treated cells, ii) identify sub-populations of cells with heterogeneous response to the treatment, and iii) reveal differential co-regulation of histone post-translational modifications in the context of drug treatment. The schPTM method enables comprehensive investigation of chromatin heterogeneity at single-cell resolution and provides further understanding of the histone code.

## Introduction

Epigenetic states are imprinted through chemical modifications to DNA and associated histone proteins that describe the complex relationship between the epigenome and cellular phenotypes^1^. The ‘histone-code’ describes the complex interplay between diverse post-translational modifications (PTMs) of histones^2^. The most prevalent histone PTMs (hPTMs) encompass acetylation (Ac), methylation (Me), ubiquitination (Ub), and phosphorylation (Ph). These hPTMs influence gene expression by modulating chromatin structure through their distinct chemical properties and their capacity to attract chromatin-modifying enzymes and binding proteins. For instance, histone 4 lysine 16 acetylation (H4K16ac) contributes to chromatin decondensation and thereby increases gene transcription^3^. In contrast, trimethylation of histone 3 lysine 27 (H3K27me3) is generally associated with transcriptional repression and has been implicated in epigenetic reprogramming^4,5^. Notably, loss of H4K16ac and H4K20me3 are common hallmarks of cancer, while several other hPTMs have been linked to neurodegenerative, autoimmune and other diseases^6–9^.

Bulk analysis of hPTMs, in which histones are extracted from cells at high purity, has markedly improved our understanding of their pivotal role in health and disease^5,10–13^. However, these bulk methods often require substantial input material (typically 1 million cells), which limits their use in cases of low or single-cell sample input (e.g. clinical specimens)^14,15^. Established single-cell profiling methods, such as single-cell RNA-sequencing, have revealed that conventional analyses of bulk samples often mask the presence of different cell types, as well as the variability of cells within the same type. Thus, single-cell analysis has emerged as a powerful tool for uncovering critical insights across diverse biomedical fields, including the identification of fibroblast clusters associated with cancer immunotherapy resistance, stem-cell programs in metastatic breast cancer, transcriptomic reprogramming in aging cardiovascular endothelial cells, and increased epigenetic variations with aging, thereby providing unprecedented resolution in understanding complex biological processes and disease mechanisms^16–19^.

While several methods have been introduced to analyze hPTMs from single cells, such as single-cell chromatin immunoprecipitation sequencing, these have had limited use in the field due sample throughput, data sparsity, technical noise, and lack of hPTM multiplexing^20–23^. Recently, Cheung et al. developed EpiTOF, for simultaneous profiling of 60 epigenetic targets in a single human immune cell using heavy metal isotope-labeled antibodies and mass cytometry by time-of-flight^5^. Interestingly, this method was able to provide insights into age-related changes in chromatin by demonstrating an increase in hPTM cell-to-cell variability during immune cell aging.

However, a fundamental limitation of the aforementioned approaches is that they rely on antibodies, which suffer from i) cross-reactivity, ii) steric hindrance when targeting combinatorial hPTMs, iii) limited number of targets, iv) quantitative ambiguity resulting from differences in affinity, v) lack of multiplexing, and vi) limited detection of novel modifications^24–26^. In contrast, liquid chromatography coupled with tandem mass spectrometry (LC-MS/MS) provides a comprehensive view of the histone code through the untargeted and quantitative measurement of a multitude hPTMs^15,27–32^. Recent advances in MS instrumentation have greatly improved sensitivity, quantitative accuracy, and throughput, making LC-MS/MS the gold standard for global single-cell proteomics^33–35^. Although histones are commonly identified in such workflows due to their abundance within the cell, a robust method for accurate quantification of hPTMs at single-cell resolution is currently lacking.

Here, we present a method that enables robust detection and quantification of single-cell histone post-translational modifications (schPTM) across the canonical histones H1, H2A, H2B, H3 and H4. We combined single-cell dispensing and semi-automated sample preparation using a cellenONE instrument coupled with a Bruker timsTOF Ultra mass spectrometer. Our schPTM workflow was able to identify 68 histone peptidoforms, comprised of 25 unique PTMs, in a single-cell. We demonstrated high quantitative accuracy and precision of our single-cell data through comparisons with bulk samples, previous data, and titration curves. Importantly, we propose a strategy to distinguish technical noise from biological noise (cell-to-cell variation) using histone standards. Finally, we treated cells *in vitro* with a histone deacetylase (HDAC) inhibitor to perturb hPTMs and identified cellular heterogeneity and differential hPTM co-regulation in response to the treatment.

This schPTM method represents a significant leap forward in single-cell epigenomics, complementing existing single-cell genomics and transcriptomics approaches. By enabling unbiased, multiplexed quantification of hPTMs at single-cell resolution, schPTM fills a critical gap in our ability to study chromatin states in heterogeneous cell populations. This method has the potential to uncover novel insights into epigenetic regulation of various biological processes, including cellular differentiation, disease progression, and response to therapeutic interventions. In the following sections, we detail the development, validation, and application of schPTM, demonstrating its robustness and utility in advancing our understanding of the histone code.

## Results

### Identification of hPTMs at single-cell resolution using LC-MS/MS

To analyze hPTMs with bulk methods, histones are extracted from millions of cells using sulfuric acid followed by trichloroacetic acid (TCA) precipitation^10,11,31,36^. Due to the high content of lysine residues in histone proteins, extracted histones are subsequently derivatized with propionic anhydride at the protein and peptide level to modify N-termini and free lysine residues. Propionylation is critical to hPTM analysis by LC-MS/MS as it ensures a sufficient size and hydrophobicity of histone peptides required for LC-MS/MS analysis. Recently, automated protocols have demonstrated that these steps can be automated using liquid handling, which allowed for increased throughput when profiling hPTMs of chemical perturbations across multiple biological backgrounds^32^. Our schPTM protocol builds off previous single-cell proteomics protocols by utilizing the cellenONE (Cellenion), a robotic picoliter dispensing platform, to automate the sample preparation steps of single-cell isolation, cell lysis, propionylation, and digestion^34,35,37,38.^

The experimental workflow, depicted in **Fig. 1A**, illustrates the direct isolation of single cells to a 384-well plate (**Supplementary Figs. 1A-D)** pre-filled with 1 µL cell lysis buffer, consisting of 0.2% (w/v) n-Dodecyl β-D-maltoside (DDM) in 1M Triethylammonium bicarbonate (TEAB)^39^. We implemented 384-well plates due to their reduced well diameter and depth relative to 96-well plates. After each step, the plate was sealed with adhesive aluminum foil, vortexed for 5 sec, and centrifuged at 500 g for 30 sec to ensure thorough mixing. To minimize evaporation or incomplete sealing of the plate, the outermost wells were excluded from sample processing. Following cell lysis, histone proteins were propionylated by dispensing 100 nl of a 25% propionic anhydride and 75% acetonitrile solution followed by incubation for 30 min at room temperature (**Supplementary Figs. 1E-G)**. Of note, dispensing of only 100 nL low-viscosity derivatization reagent with the cellenONE required fine-tuning of the piezo dispensing capillary nozzle parameters to obtain a stable droplet (**Supplementary Fig. 1E; Methods**). Next, the plate seal was removed and the temperature of the cellenONE plate holder was increased to 50°C, ensuring complete evaporation of the propionylation reagents which removes any chemical interference during the next digestion step. Following this, 1 µl digestion buffer is added, consisting of 3 ng/µl trypsin, 10 U/µL Benzonase, and 1% ProteaseMAX in 1M TEAB. The plate is then sealed and incubated for 2 hrs at 50°C within the cellenONE for histone digestion. The free amino termini of the digested peptides were then propionylated as described above to improve their chromatographic retention. The dual propionylation steps, on average, effectively derivatized 77% of all unique peptides (histone and non-histone peptides) identified in single cells (**Supplementary Figs. 2A-D; Methods)**. Finally, samples were manually diluted with 0.1% TFA and stored at -80 °C prior to LC-MS/MS analysis. To serve as a reference and assess the accuracy of our workflow, we also prepared samples consisting of ∼100 cells (100-cell bulk), which were processed identically to the single cells (**Methods**).

**Fig. 1.**
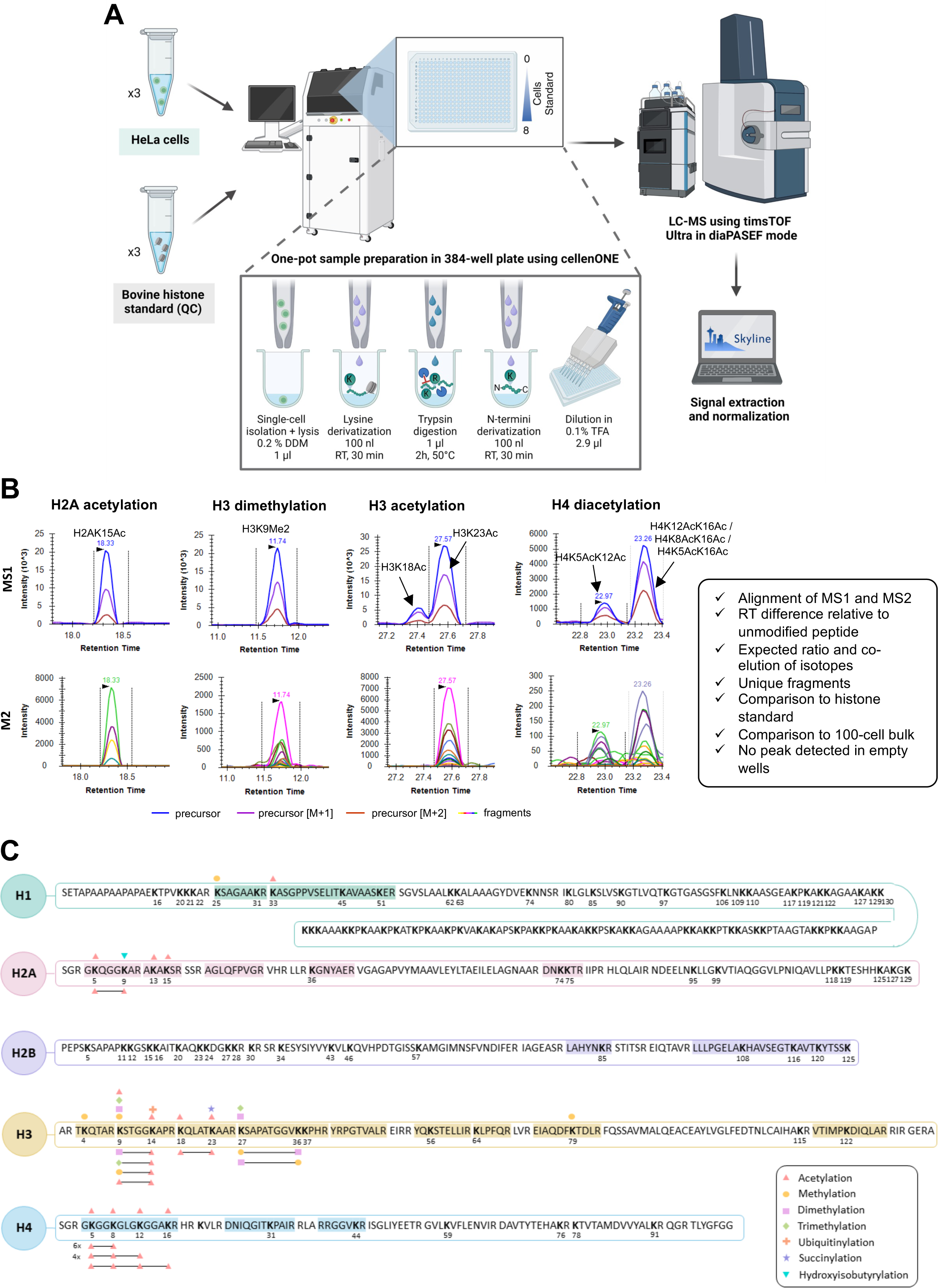
Overview of schPTM method and identified peptidoforms. **A)** Schematic overview of the method, from left-to-right: 1) Suspensions of single cells or histone standards were isolated and prepared within the wells of a 384-well plate using the liquid handling capabilities of the cellenONE instrument, 2) Histone peptidoforms were acquired using the Vanquish LC and TimsTof Ultra MS instruments, 3) Data analysis was performed with Skyline and custom R scripts. For details on cellenONE cell isolation and preparation see **Supplementary Fig. 1**. **B)** Extracted ion chromatograms (XICs) of MS1 precursor (top panels) and MS2 fragments (bottom panels) for four representative histone peptidoforms in a representative single-cell. Peaks are picked based on criteria shown in box on right side. For additional XICs of isobaric peptides see **Supplementary Fig. 3A**. **C)** Overview of the histone peptidoforms identified by the schPTM method. Detected peptide sequences are highlighted. PTMs are represented by symbols shown in the legend on the bottom-right. Single PTMs are shown on top of sequences, while combinatorial PTMs are shown by a connecting line underneath the sequence. For proportions of each modification see **Extended Data Fig. 1A**.

Resulting MS data were analyzed in Skyline^40^ using a spectral library derived from 100-cell bulk samples, with manual adjustment of peak boundaries based on predetermined requirements (**Fig. 1B**). For confident peptide identification, we used a combination of filtering criteria: 1) clear distinction of peptide signal versus measurement noise, 2) MS1 and MS2 signal alignment, 3) expected ratio and co-elution of isotopes, 4) the delta in retention time to the unmodified peptide, 5) the presence of unique fragment ions indicative of a particular PTM, 6) comparison to 100-cell bulk samples, and 7) comparison to histone standards (**Fig. 1B, right box**).

Following all these criteria, we identified peptides from H1, H2A, H2B, H3, and H4, as well as their variants H1.2, H1.5, H2AJ, H2AX, and H3.3 (**Supplementary Table 1**). We were able to identify peptidoforms that were acetylated (Ac), methylated (Me), dimethylated (Me2), trimethylated (Me3), ubiquitinylated (Ub), succinylated (Su) and hydroxyisobutyrylated (Hib) (**Extended Data Fig. 1A**). These were present either individually (25 peptidoforms) or in combination with other PTMs (43 peptidoforms), resulting in 68 distinct peptidoforms (**Fig. 1C**). Notably, we achieved comprehensive coverage of the N-terminal tails of H3 and H4, in particular of peptides H3(9-17) and H4(4-17) that were identified with multiple peptidoforms. Missing regions included the H2B N-terminus and H1 C-terminus which lack arginine residues, resulting in long and hydrophobic peptides. Identification of these peptides is challenging for LC-MS/MS analysis owing to their poor chromatographic behavior and the presence of several peptidoforms sharing the same precursor mass.

### Benchmarking of schPTM quantification

To benchmark quantitative accuracy of our method, we prepared 3 batches of HeLa cells on separate days (**Supplementary Table 2)**, totaling 60 samples with varying amounts of cells as input (**Extended Data Fig. 1B**). A batch is defined as a single 384-well plate that includes single-cell and control samples (i.e. cell and bovine histone standard titration curves) that were prepared and acquired together. We consistently detected 68 histone peptides across all samples with < 0.1% missing values (**Extended Data Fig. 1C**), highlighting the strength of the data independent acquisition (DIA) method. To distinguish between a positive peptidoform signal and measurement noise, we compared the raw intensities of single-cell samples to ‘empty wells’, which were samples that contained only reagents and no cell. Despite the expectedly low signal from single cells, we found that the MS1 signal intensities in the single-cell samples were on average ∼9.5X above the empty wells (i.e. 0 cells; **Fig. 2A** and **Extended Data Figs. 2A-B**). Peptides from single cells that had similar MS1 signal intensities in ‘empty wells’ were considered as background noise and excluded in subsequent analyses (**Fig. 2B** and **Extended Data Fig. 2C; Methods**). Moreover, MS1 intensities in our measurements were on average over 3X greater than MS2 intensities and generally suffered less from distortion at lower signal intensities **(Extended Data Fig. 1D**). We therefore quantified histone peptide abundances at the MS1 level, similarly to MS-based histone analysis using EpiProfile^41^. Exceptions to this were co-eluting isobaric modifications of the H4(4-17) histone peptide, which contains four modifiable lysine residues and is often present in several isobaric forms due to different degrees of acetylation. To deconvolve these signals for relative quantification of each isoform, we used the MS2 fragment ion intensities indicative of the modification or combinations thereof for each peptidoform as shown in **Supplementary Figs. 3A-B** and detailed in **Methods**.

**Fig. 2.**
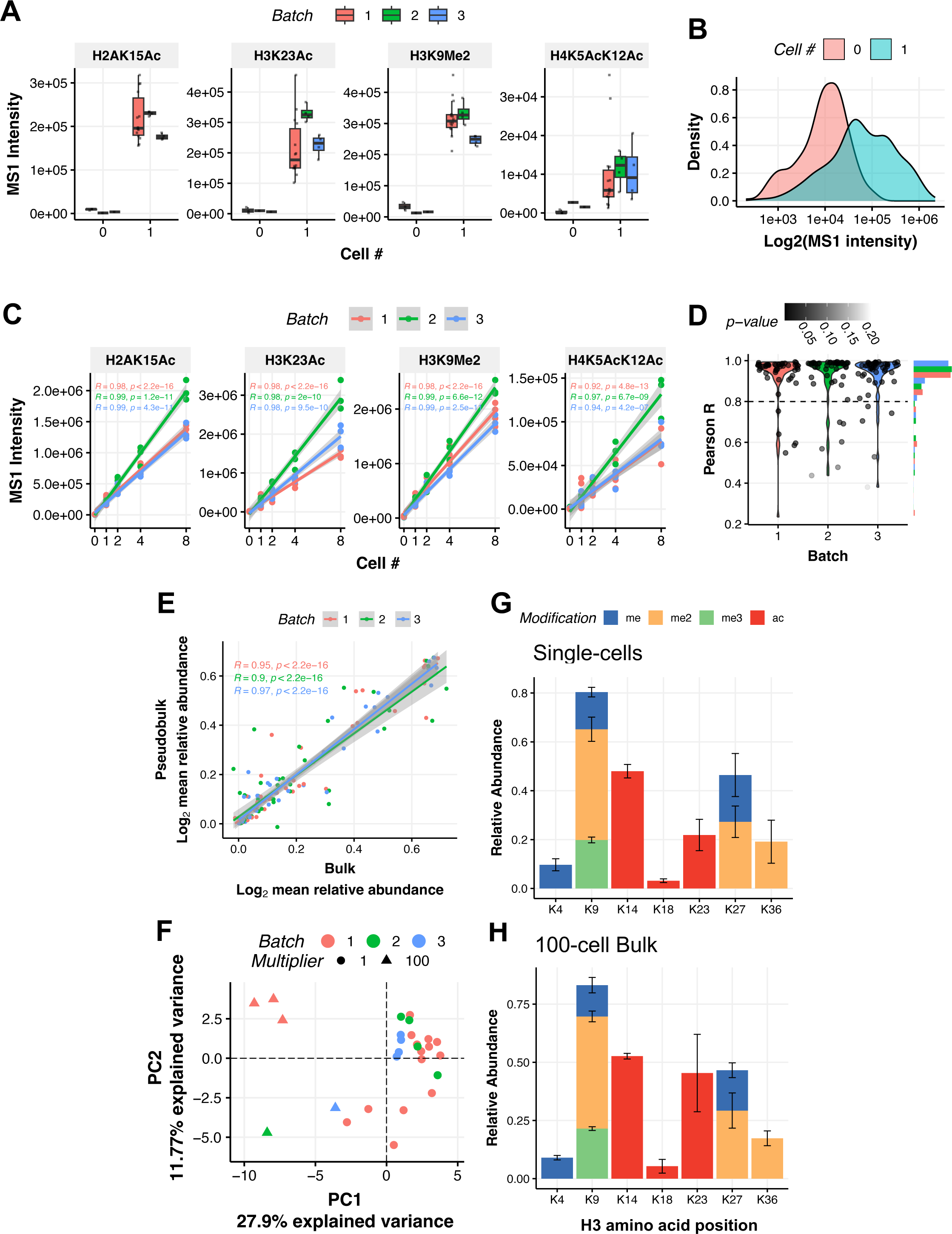
schPTM quantification enabled by high sensitivity and precision. **A)** Comparison of raw MS1 intensity of empty wells (Cell #0) and wells containing single cells (Cell # 1) for 4 representative histone peptidoforms for each batch. For all peptidoforms see **Extended Data Fig. 2A**. **B)** Distribution of raw Log_2_ MS1 intensities for all histone peptidoforms in batches 1-3 for empty wells and wells containing single cells after filtering out peptides with low signal (**Methods**). For each batch separately see **Extended Data Figs. 2B-C**. **C)** Titration curves for 4 representative histone peptidoforms of raw MS1 intensity where various amounts of cells were prepared in a single-well and then were assessed for linearity. Statistics: Significance of Pearson R correlation coefficients for each batch was tested using a t-test. For all peptides see **Extended Data Fig. 3B**. **D)** Distribution of Pearson R correlation coefficients for titration curves of all histone peptidoforms in each batch. A dashed line is drawn at R=0.8 to indicate the threshold used for filtering out histone peptidoforms deemed to be non-quantifiable. Statistics: Significance of Pearson R correlation coefficients for each batch was tested using a t-test. **E)** Correlation of histone peptidoform Log_2_ relative abundance between bulk (∼100-cell) and ‘pseudobulk’ (mean of single cells) samples in baches 1-3. Statistics: Significance of Pearson R correlation coefficients for each batch was tested using a t-test. For correlation using raw MS1 intensity see **Extended Data Fig. 6A**. **F)** Principal component analysis of histone peptidoform relative abundances for single-cell and 100-cell bulk samples. For no batch-correction see **Extended Data Fig. 5A-C**. For direct comparison of single-cell and bulk samples see **Extended Data Fig. 6B-C**. **G)** Individual histone PTMs (single and combinatorial PTMs) relative abundances detected on histone H3 in single cells and **H)** 100-cell bulk samples from batches 1-3. For comparison to previous bulk studies, see **Extended Data Fig. 6D-E**.

To benchmark the quantitative accuracy of our method, we assessed the linearity of signal intensities using a 5-point titration curve of varying cell inputs (i.e. 0, 1, 2, 4, and 8 cells) and a commercially available bovine histone standard from calf thymus (Roche) (0, 10, 20, 40, and 80 pg; 1 cell contains ∼6.5 pg of histone protein^42^) prepared with our schPTM workflow. As expected, the MS1 intensity was proportional to the number of cells and expected bovine histone standard across almost all peptidoforms yielding Pearson R values consistently > 0.9, demonstrating quantitative accuracy of the method (**Figs. 2C-D**, **Extended Data Figs. 3A-B and 4A-C**). Based on this assessment, peptides with an average R value of less than 0.8 were excluded from subsequent analysis to ensure accurate quantification (**Extended Data Fig. 3B; Methods**).

To further assess the accuracy of hPTM quantification in single cells, we compared the 100-cell bulk samples to single cells. After quality control filtering, normalization, and batch-correction (**Extended Data Figs. 5A-F; Methods**), we were able to quantitatively analyze 55 peptidoforms across 5 100-cell bulk samples and 22 single cells (**Extended Data Fig. 1B**). Data were normalized using the peptide ratio method as previously described^41^, followed by a Log(1 + *x*) transformation. We averaged the peptidoform relative abundances across all single cells within each batch to create a cumulative ‘pseudobulk’ sample. We then correlated the ‘pseudobulk’ peptidoform relative abundances to the average of the bulk samples within each batch which yielded an average R value of 0.94 (**Fig. 2E; Methods**). Similar results were also obtained without normalization (average R value across 3 batches: 0.89; **Extended Data Fig. 6A**). The high correlation between the single-cell pseudobulk and 100-cell bulk samples indicates accurate quantification, while the variation seen in this correspondence is indicative of the single-cell precision of our data (**Fig. 2E**).

We then performed principal component analysis (PCA) on the relative abundances of all peptidoforms. The single-cell and 100-cell bulk samples separated along the first principal component (PC) and accounted for 27.9% of the variance in the data (**Fig. 2F**). This difference is not due to large systematic differences as the average difference of hPTM relative abundances between single-cell and the mean of 100-cell bulk samples was only 1.28×10^-3^ Log_2_-fold (**Extended Data Fig. 6B**). Nonetheless, some small, but consistent, differences can be noted, such as higher relative abundance of H3K9me2 and lower relative abundance of H3K23ac in single cells as compared to 100-cell bulk samples (**Extended Data Fig. 6C**). As the signal of peptidoforms with such differences were shown to be sufficiently above background levels (**Extended Data Fig. 2A**) and have high quantitative accuracy (**Extended Data Fig. 3B**), the observed differences are suggestive of higher accuracy of the schPTM method compared to bulk samples with larger input, which may be due to less interference during sample preparation and acquisition. Finally, we compared the relative abundances of individual PTMs on histone H3 between single cells, 100-cell bulk samples, and bulk HeLa cell samples from previous studies where we found high correspondence of single-cell with bulk samples (**Figs. 2G-H; Extended Data Figs. 6D-E**). Collectively, we were able to validate the accuracy of our schPTM quantification against traditional bulk methods.

### Biological cell-to-cell variability of hPTMs

To distinguish between technical and biological variability in single-cell samples, we first used technical replicates of a commercially available bovine histone standard titrated to various amounts (as described in the previous section). This was made possible due to the highly accurate and precise picoliter dispensing of the cellenONE instrument. After quality control filtering, normalization, and batch-correction (**Methods**), we determined the coefficient of variation (CV) of the relative abundances of 55 peptidoforms prepared within a batch (intra-) or across batches (inter-). For the 10 pg histone standard samples (roughly equivalent to the amount of histone in a single-cell), both intra- and inter-CVs were below 25% on average (**Figs. 3A-B**). For data that were unnormalized and not batch-corrected, intra-CVs were still below 25%, while inter-CVs were roughly double, demonstrating the impact of normalization and batch-correction (**Extended Data Figs. 7A-B**). Moreover, we compared the 10 pg histone standard samples to technical replicates of bulk HeLa cells^43,44^ and processing replicates of cell lines treated with DMSO in the NIH Library of Integrated Network-Based Cellular Signatures consortium (LINCS)^32^ (**Supplementary Table 3).** With this, we found that hPTM CVs were comparable to the bulk HeLa data^43,44^ (**Fig. 3C, left and middle panels**), as well as the cell lines from the LINCS dataset^32^ (**Fig. 3C, right panel**). Overall, this analysis established a baseline for technical variation expected from single-cell sample input amounts within a batch (intra-) or across batches (inter-) for our schPTM workflow, highlighting the technical reproducibility of our schPTM method.

**Fig. 3.**
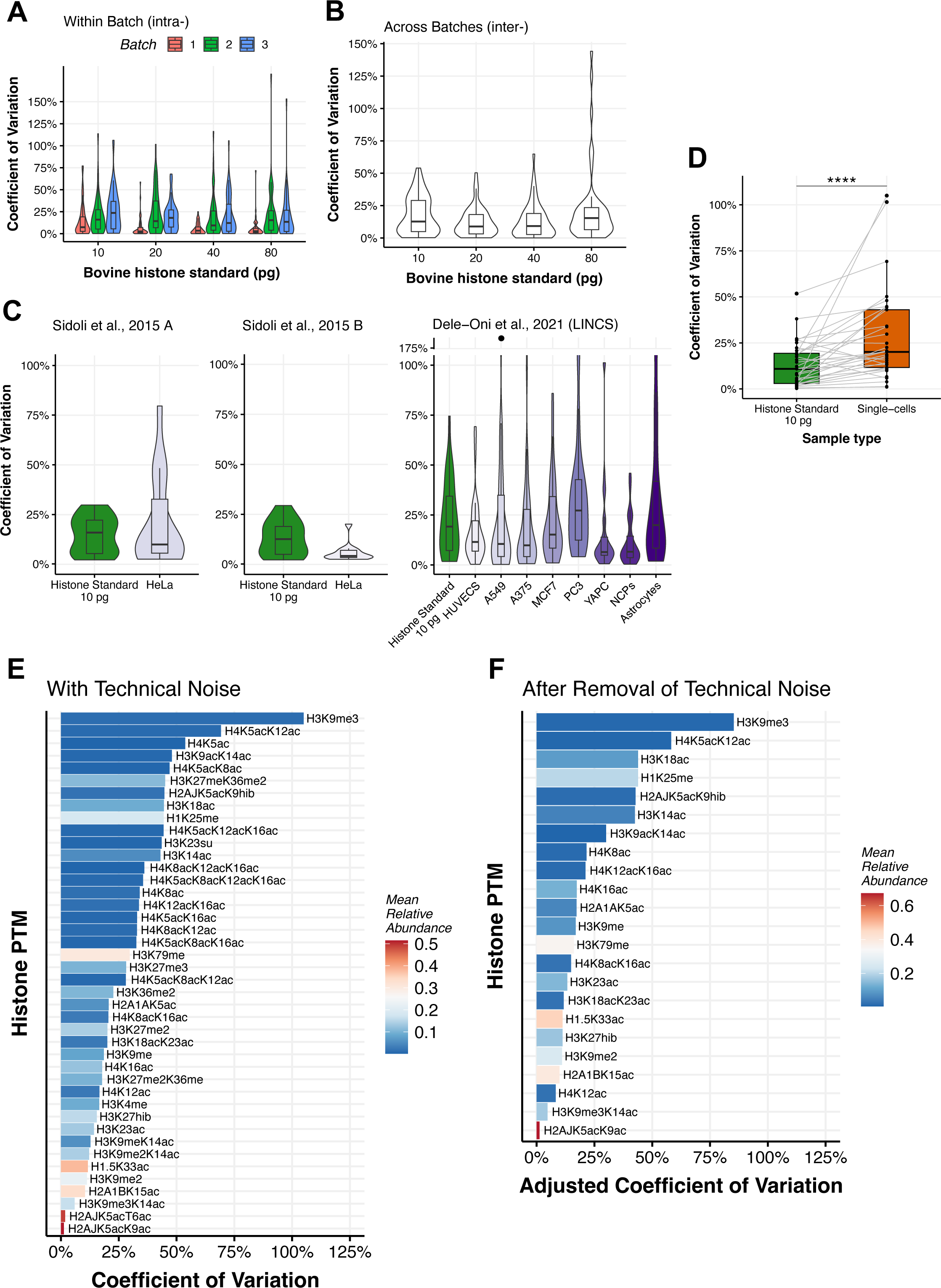
Low technical noise allows for analysis of hPTM biological cell-to-cell variability. **A)** Coefficient of variation (CV) of histone peptidoform relative abundances for histone standard technical replicates within each batch (intra-) and **B)** across the batches 1-3 (inter). For results using raw MS1 intensities see **Extended Data Figs. 7A-B**. **C)** CV of histone peptidoform relative abundances for 10 pg histone standard technical replicates across batches 1-3 compared to bulk technical replicates of HeLa cells (Sidoli et al., 2015 A, 5 replicates, 16 peptidoforms; Sidoli et al., 2015 B, 4 replicates, 12 peptidoforms) and processing replicates of cell lines treated with DMSO in the NIH Library of Integrated Network-Based Cellular Signatures Consortium (Dele-Oni et al., 2021, 3 replicates, 29 peptidoforms). Histone peptidoforms were filtered to match those quantified in each respective study (**Supplementary Table 3**). **D)** CV of histone peptidoform relative abundances for 10 pg histone standard across batches 1-3 compared to that of single cells and matched by peptidoform. Statistics: paired Wilcox test, p-value=1.1*10^-5^ (****). **E)** CV of histone peptidoform relative abundances of peptidoforms ranked from highest to lowest amongst single-cells from batches 1-3 shown with technical noise and **F)** after removal of technical noise (i.e. cell-to-cell noise).

An important capability for single-cell analysis is to be able to quantify subtle cell-to-cell biological variation that can reveal cell states and regulatory relationships inaccessible by bulk analyses. Indeed, our method allows for this possibility as the CVs of peptidoforms across single cells were significantly higher than those amongst the technical replicates (**Fig. 3D**). To then determine biological cell-to-cell variability for individual peptidoforms, we adjusted for technical noise (determined by the 10 pg histone standard samples) and ranked peptidoforms by their CV (**Figs. 3E-F; Extended Data Fig. 7C; Methods**). Importantly, this analysis revealed H3K9me3 and H4K5acK12ac as the most biologically variable peptidoforms in our cell model, likely due to variation in cell cycle phase, whereas H3K9me3K14ac and H2AJK5acK9ac were found to be least variable within the analyzed cell population (**Fig. 3F**).

### Distinct single-cell hPTM profiles following HDAC inhibitor treatment

To determine if our schPTM method can detect the presence and change in abundance of biologically relevant hPTMs due to chemical perturbation, we treated HeLa cells with 5 mM of the histone deacetylase (HDAC) inhibitor sodium butyrate (NaBut). NaBut competitively binds to the zinc sites of class I and II HDACs, resulting in hyperacetylation of several histone sites, particularly those located on histones H3 and H4. After a 24-hour incubation, both control (PBS vehicle) and NaBut treated cells were harvested and processed using our schPTM workflow in 2 batches (**Fig. 1A**). Single cells were isolated from three different cell populations, control, treated, and a 1:1 mixture of control and treated (mixed) (**Fig. 4A; Supplementary Table 4)**. We included a mixed population of cells to evaluate if these cells can be classified based on their hPTMs profiles without prior knowledge of the control or treated groups. Like previous experiments, we also included 100-cell bulk samples. Following our quality control filtering, normalization, and batch-correction pipeline (**Methods**), we were able to confidently quantify 53 peptidoforms in 5 controls, 12 treated, and 33 mixed single cells (**Extended Figs. 8A-D**). Due to the low number of control cells that passed filtering criteria in this experiment, we further included 22 untreated single-cells and 5 100-cell bulk samples from the benchmarking experiments in the analysis (**Extended Data Fig. 8E)**. PCA of all analyzed samples revealed clear separation between control and treated cells in PC1, accounting for 27.89% of the variation, which was significantly correlated to the NaBut treatment variable (R=0.66, p<0.001). Mixed cells were uniformly distributed across PC1, as expected. Separation in PC2 was mainly driven by differences in single-cell and 100-cell bulk samples (R=0.81, p<0.001). (**Fig. 4B)**.

**Fig. 4.**
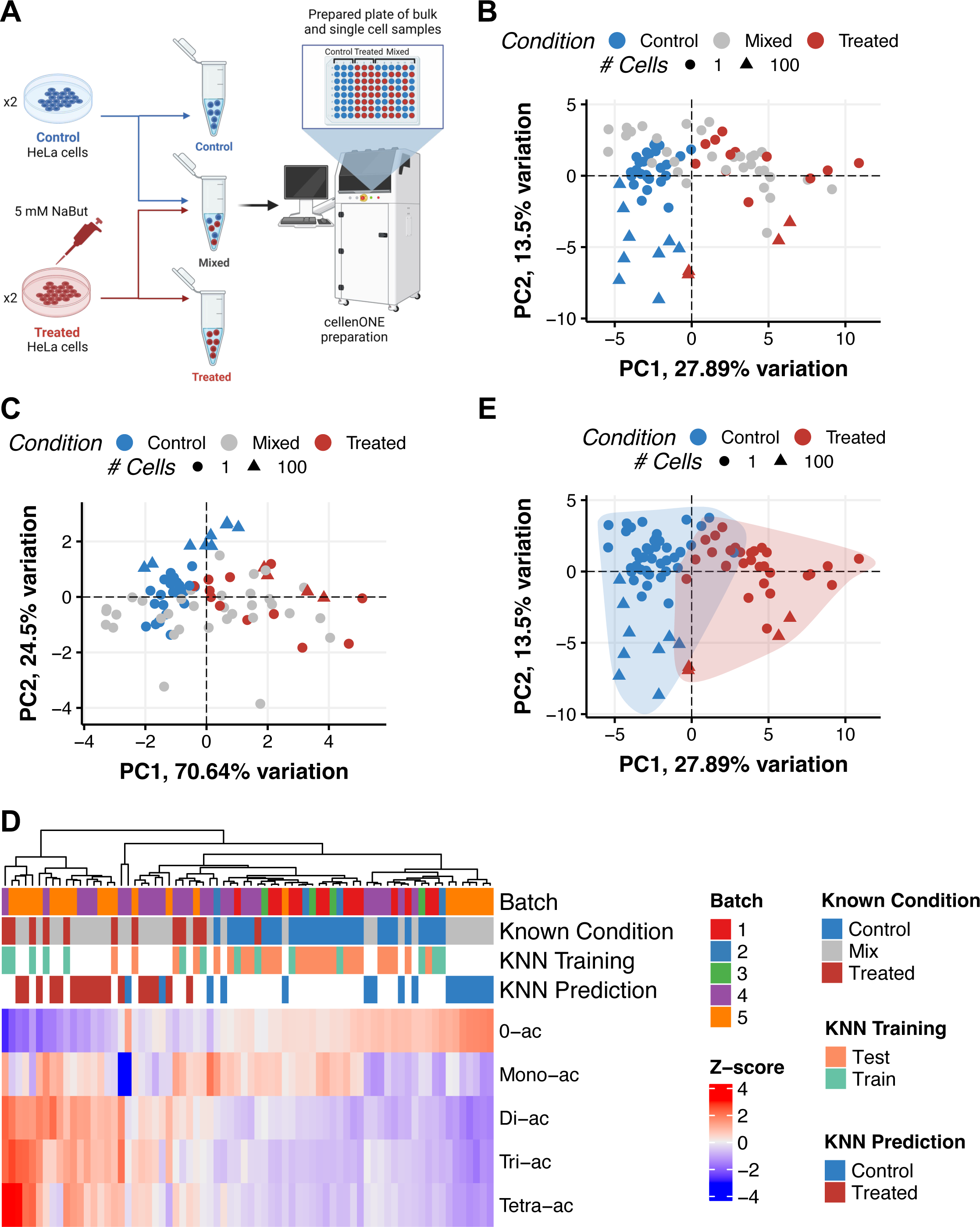
Classification of single cells following HDAC inhibitor treatment. **A)** Schematic overview of experimental design where after HDAC inhibitor treatment, control and treated cell suspensions are also mixed 1:1 to create the ‘mixed’ group where the label of each cell was unknown. For plate layout see **Supplementary Table 4**. **B)** Principal Component Analysis (PCA) of histone peptidoform relative abundances for single-cell and 100-cell bulk samples from HDAC inhibitor experiment. Note that samples from benchmarking experiments were included to increase number of control cells, for details see **Extended Data Fig. 8E**. **C)** PCA of same samples in (**B**), but instead calculated using summarized relative abundances of H4 peptide acetylation (i.e. mono, di, tri and tetra) which were used for classifying the mixed cells into control and treated groups as shown in (**D**). **D)** Heatmap and hierarchal clustering of scaled H4 peptide acetylation summarized relative abundances (Z-score) for single cells as shown in (**C**). Cells are shown in columns and scaled H4 peptide acetylation types are shown in rows. A description of the annotation rows at the top of the heatmap (also shown in legend) starting from the 1^st^ row and moving down to the 4^th^ row is as follows: 1^st^ row) Batch each cell was prepared in; 2^nd^ row) Known group of cells used to train and test the k-nearest neighbors (KNN) classifier, as well as validate the accuracy of the predictions; 3^rd^ row) Random assignment of cells with known group (control or treated) to training (50%) and test (50%) groups to assess KNN classification accuracy; 4^th^ row) Predicted labels of ‘mixed’ cells using KNN classification after training using all cells with known group. **E)** PCA, as shown in (**B**), with ‘mixed’ cells now classified as control or treated as predicted by KNN classification as shown in (**D**).

To classify the mixed cells, we calculated the relative abundance of the different acetylations on histone H4 (amino acids 1-16; **Methods**), known to be regulated by NaBut treatment^13^. PCA of relative abundances of mono, di, tri and tetra-H4 acetylation clearly separated the control and treated cells along PC1 and was significantly correlated to the NaBut treatment variable (R=0.54, p<0.001) (**Fig. 4C**). Importantly, focusing on only H4 acetylation patterns increased the explained variance of PC1 to 70.64%, compared to 27.89% when analyzing all peptidoforms (**Fig. 4B**). The informed selection of the classifying histone modification subset was therefore used to train a k-nearest neighbor (KNN) classifier on the ‘known’ cells, which achieved an accuracy of 84.21% in our evaluations (**Methods**). We then used this classifier to assign labels of ‘control’ or ‘treated’ to 16 or 17 single cells from the mixed population, respectively (**Fig. 4D-E**). This demonstrates the ability of our method to classify cells using informed selection of hPTM subgroups without knowing the assigned condition *a priori*. This is particularly critical in the analysis of complex cell populations from tissue that contain different cell types or different cellular states at distinct ratios.

### HDAC inhibitor treatment induces histone hyperacetylation and cell-to-cell chromatin heterogeneity

We next performed differential abundance analysis to assess changes in the abundance of all peptidoforms using 43 control and 29 treated single cells (**Methods)**. As expected, we found the majority of the acetylated peptidoforms of H1.2, H2A.1, H3 and H4 to be significantly increased in the treated group relative to the controls (**Fig. 5A; Supplementary Table 5**). The largest increase in relative abundance was observed for the tri-acetylated H4 peptidoforms, H4K5acK12acK16ac, H4K8acK12acK16ac, and H4K5acK8acK16ac. In support of this, analysis of relative abundance of the different acetylations on histone H4 revealed a significant increase in di-, tri- and tetra-acetylated hPTMs independent of the mono-acetylated hPTMs (**Fig. 5B**). Analysis of global relative abundances of each modification type (i.e. acetylation, mono-methylation, di-methylation and tri-methylation), confirmed the significant increase in acetylation, while also revealing a corresponding decrease in methylation (**Fig. 5C**). The results of this analysis were supported by the same analysis using 100-cell bulk samples (**Extended Data Figs. 9A-C; Supplementary Table 6**).

**Fig. 5.**
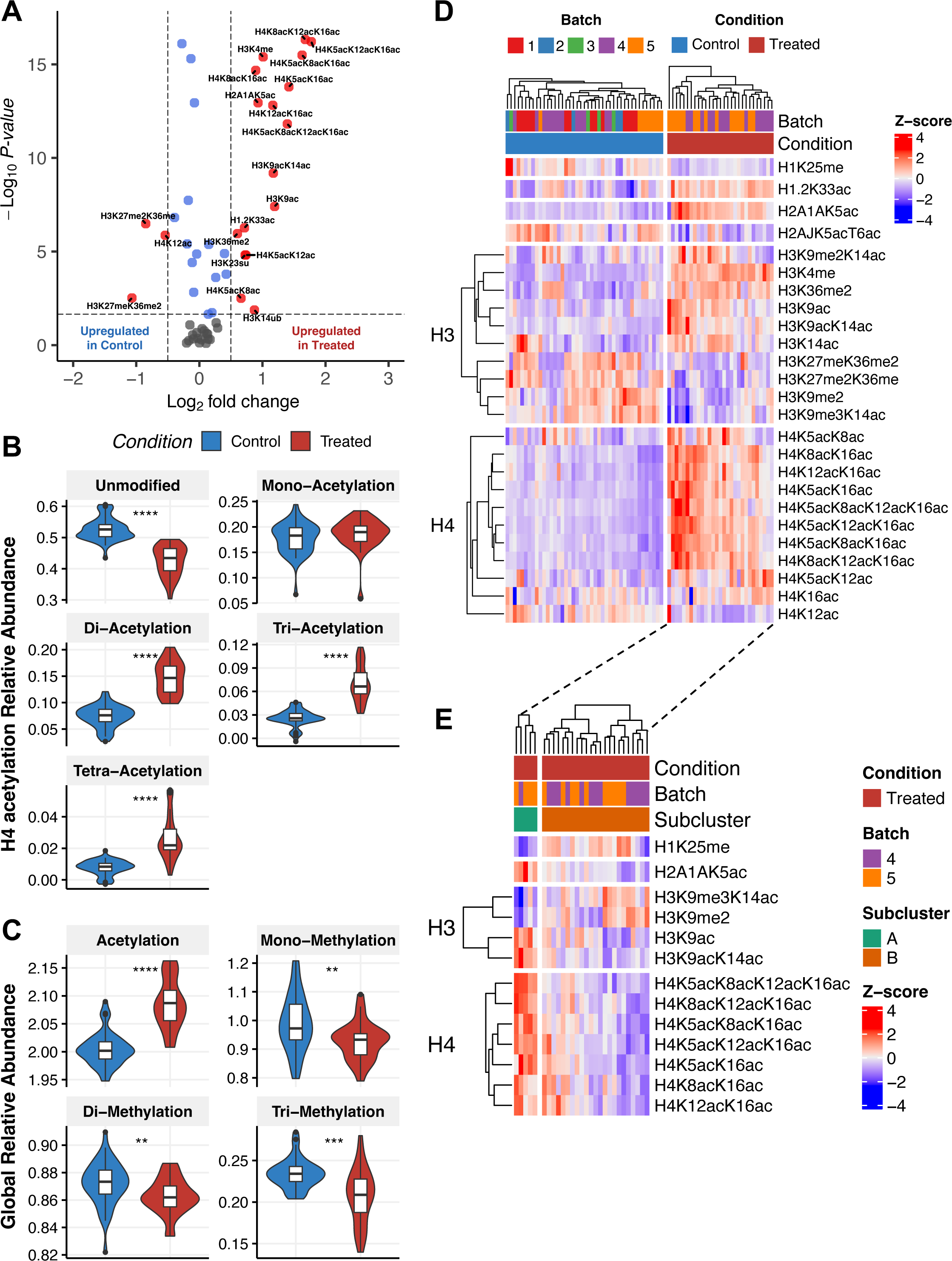
HDAC inhibitor treatment induces histone hyperacetylation and cell-to-cell chromatin heterogeneity. **A)** Differential abundance of modified histone peptidoforms relative abundances in single cells for treated vs. control comparison. Statistics: Wilcox test with Benjamini & Hochberg multiple testing correction. Red points are peptidoforms that are significant (adjusted p-value < 0.05) and Log_2_ fold change > 0.5. Blue points are peptidoforms that are significant only. Grey points are peptidoforms that are not significant (adjusted p-value > 0.05) and Log_2_ fold change < 0.5. For table of full results, see **Supplementary Table 6**. For results of bulk samples, see **Extended Data Fig. 9** and **Supplementary Table 7**. **B)** Summarized relative abundances of global PTMs in single cells for control and treated groups. Statistics: Wilcox test, ** p <= 0.01, *** p <= 0.001, **** p <= 0.0001. **C)** Summarized relative abundances of H4 peptide acetylation in single cells for control and treated groups. Statistics: Wilcox test, ** p <= 0.01, *** p <= 0.001, **** p <= 0.0001. **D)** Heatmap and hierarchal clustering of modified histone peptidoform scaled relative abundances (Z-score) that were significantly differentially abundant (adjusted p-value < 0.05) between treated vs. control groups as shown in (**A**). Cells are shown in columns and histone peptidoforms are shown in rows. Peptidoforms are grouped by histone proteins (left side). **E)** Sub-clustering results of treated group as shown in (**D, right side**), showing 2 subclusters (A & B) of cells identified within the treated group and the modified histone peptidoforms that are significantly differentially abundant (adjusted p-value < 0.1) between the subclusters. Subclusters were identified at the 2^nd^ level of the hierarchal clustering tree shown in (**D, top-right**). Statistics: Wilcox test with Benjamini & Hochberg multiple testing correction. For full subcluster differential abundance results, see **Extended Data Fig. 9E** and **Supplementary Table 8**.

Interestingly, the relative abundance of H3 and H4 peptidoforms that were found to be significantly differentially abundant between control and treated groups (**Fig. 5A, red points**), were observed to vary from cell-to-cell in the treated group, indicative of a heterogenous response to the NaBut treatment (**Fig. 5D, right side**). Thus, we sub-clustered the treatment group and performed differential abundance analysis on the 2 resulting clusters (**Fig. 5E; Extended Data Fig. 9E; Supplementary Table 7; Methods**). Subcluster A was characterized by relatively low H3Kme3K14ac and H3K9me2, and hyperacetylation of H4 tails, which are all associated with active transcription. In contrast, cells within subcluster B showed relatively high H3Kme3K14ac and H3K9me2 and lower levels of H4 acetylation, which are all associated with repressed transcription (**Fig. 5E**). Importantly, the differential response to NaBut treatment was not detected when using 100-cell bulk samples to perform the same analysis (**Extended Data Fig. 9D**). In summary, this analysis demonstrates that our schPTM method was able to recapitulate the expected effect of NaBut treatment, in addition to enabling a fine-grained analysis of subtle differences in drug responses within a relatively homogeneous population of cells *in vitro*.

### Differential hPTM crosstalk induced by HDAC inhibitor treatment revealed by co-variation analysis

The single-cell resolution of our schPTM data has the potential to enable study of the co-variation of hPTM abundances, which is most often masked by the averaging that occurs in bulk sample analysis. To perform hPTM co-variation analysis, we correlated the trends of each peptidoform relative abundance across single cells to one another and analyzed the significant correlations (**Methods**). This resulted in covariance matrices for the control (**Fig. 6A**, **lower-left triangle**; 177 significant correlations, Two-sided T-test p-value<0.05) and treated groups (**Fig. 6A**, **upper-right triangle;** 206 significant correlations, Two-sided T-test p-value<0.05), which were hierarchically clustered in reference to the control group (**Supplementary Table 8**). In the control group, a group of hPTMs, encompassing acetylations associated with open chromatin and gene expression^3^ (i.e. H4K8acK12acK16ac, H4K1212acK16ac, H4K5acK16ac, H4K8acK16ac, H4K5acK8acK16ac, H4K5acK8acK12acK16ac, and H4K5acK12acK16ac), showed significant positive correlations, as expected. In contrast, the bivalent hPTM H3K9me3K14ac, associated with a poised chromatin inactive state^45^, negatively correlated with the aforementioned acetylated peptidoforms, in addition to H2A1AK5ac, H3K9acK14ac, and H3K9me (**Fig. 6A**, **lower-left triangle**). At the global level, the covariance matrix of the treated group was significantly different from that of the control group, indicating a change in hPTM crosstalk induced by the NaBut treatment (Mantel’s test p-value=9.99*10-5). Given that an increase in the number of histone PTM-PTM correlations in the treated group may be due to an increase in cell-to-cell variability, we calculated the pairwise Euclidean distance for cells within each group. Unexpectedly, this showed a decrease in the treated group relative to the control (**Extended Data Fig. 10A)**. In addition, the hPTM CV distributions did not significantly differ between the groups (**Extended Data Fig. 10B)**. These results suggest that the observed changes in hPTM crosstalk are likely due to alterations in hPTM regulation, rather than increased overall variability in response to the HDAC treatment.

**Fig. 6.**
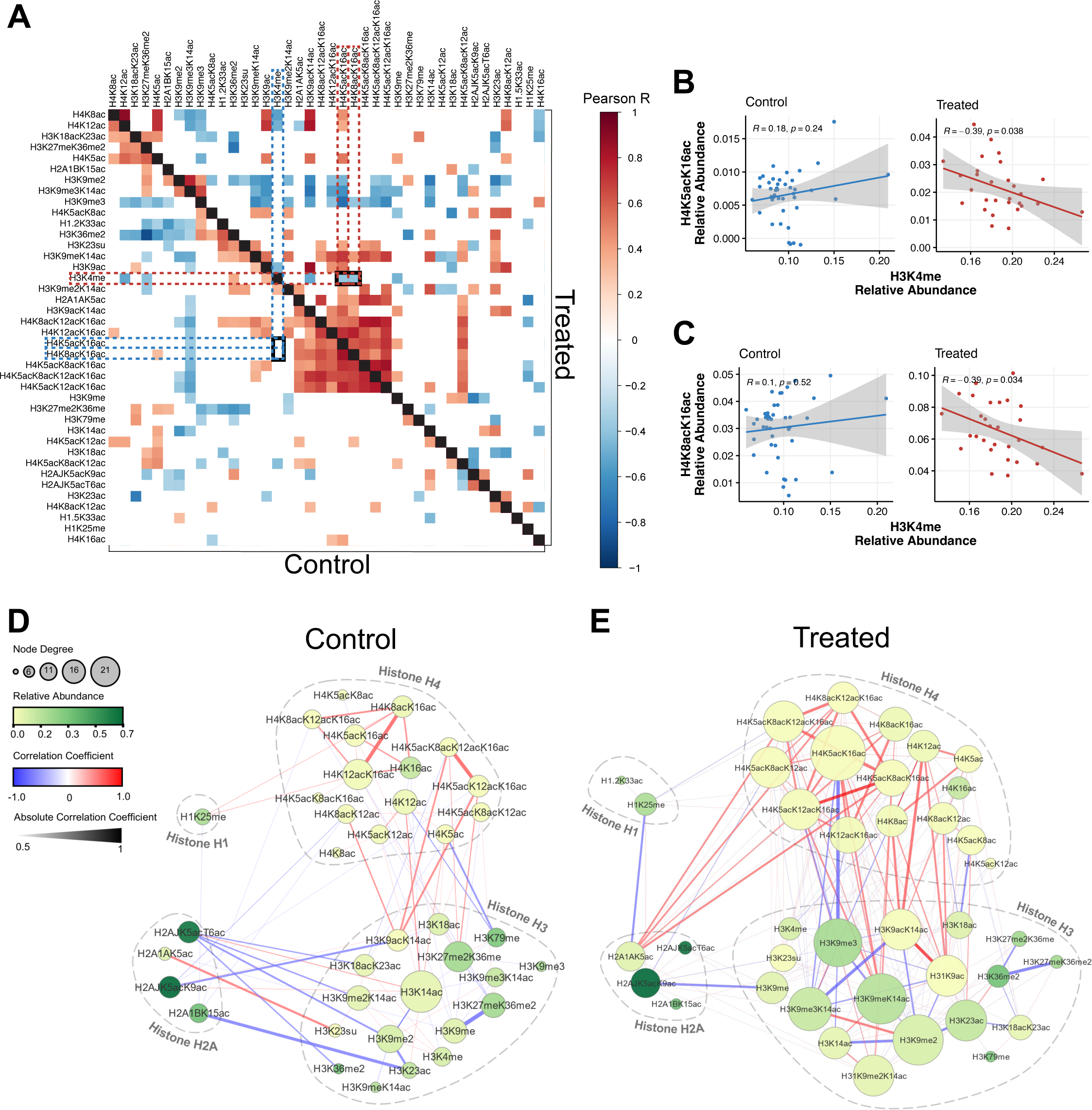
Differential hPTM crosstalk induced by HDAC inhibitor treatment revealed by co-variation analysis. **A)** Covariance matrix of modified histone PTM-PTM correlations derived from histone peptidoform relative abundances with significant Pearson R correlation coefficients (p-value < 0.05) shown as colored squares. PTM-PTM correlations for control group are shown in lower-left triangle and upper-right triangle are PTM-PTM correlations for treated group. Correlations of H4K4me with H4K8acK16ac and H4K4me with H4K5acK16ac, which are significantly altered due to NaBut treatment (adjusted p-value < 0.05), are highlighted in the control (blue dashed outline) and treated (red dashed outline) groups. Statistics: Significance of Pearson R correlation coefficients was tested using a t-test; Significance of differential Pearson R correlation coefficients was tested using a permutation test with 10,000 permutations (**Methods**). For a table of Pearson R correlation coefficients p-values, see **Supplementary Table 9**. For full results and table of differential Pearson R correlation coefficient testing between treated vs. control groups, see **Extended Data Fig. 10C-D and Supplementary Table 10**. **B-C)** Comparison of histone peptidoform relative abundances in single-cells for PTM-PTM correlations which were significantly altered due to NaBut treatment highlighted by blue (control) and red (treated) dashed lines in (**A**). Correlations of H4K4me with H4K5acK16ac are shown in (**B**) and correlations of H4K4me and H4K8acK16ac are shown in (**C**). Within each panel, the control group is shown on left sub-panel and the treated group on the right sub-panel. Statistics: Significance of Pearson R correlation coefficients for each batch was tested using a t-test. **D-E)** Network diagrams derived from covariance matrices as shown in (**A**), showing the significant Pearson R correlation coefficients (p-value < 0.05) of modified histone PTM-PTM correlations (edges) for each of the modified peptidoforms (nodes). Control group shown in (**D**) and treated in (**E**). Node size represents node degree (i.e. number of significant correlations), node color represents mean relative abundance amongst all cells in control and treated groups, edge color represents absolute correlation coefficient, and edge width represents the scaled correlation coefficient (see legend on left side).

To reveal the specific hPTMs that showed differential covariance (i.e. shifts in the co-regulation), we performed a differential analysis between the treated and control groups for each of the histone PTM-PTM correlations (**Methods; Supplementary Table 9**). Globally, there was a shift towards positive correlations in the treated group, indicating a higher degree of synchronous co-regulation in response to the NaBut treatment (**Extended Data Fig. 10C**). At the hPTM level, we observed a significant decrease in the correlations of H3K4me with H4K8acK16ac and H3K4me with H4K8acK16ac in the treated group relative to the control (**Fig. 6A, intersection of dotted lines and 6B-C; Extended Data Fig. 10D**). This was an unexpected finding, as H3K4me, H4K8acK16ac, and H4K8acK16ac were amongst the most significantly upregulated peptidoforms upon HDAC inhibitor treatment (**Fig. 5A**).

To obtain a complementary perspective of hPTM cross-talk, we visualized histone PTM-PTM correlations as networks for the control and treated groups (**Fig. 6D-E)**. This clearly revealed patterns of hPTM co-regulation both within (intra-) and across histone tails (inter). For example, in the control group, intra-H2A co-regulation were nearly absent, while inter-H2A-H3 PTM correlations were prominent, suggesting a shared regulatory mechanism (e.g. the positive correlation of H2AK5ac with H3K23su; **Fig. 6D, lower-left)**. Interestingly, highly connected hPTM hubs potentially indicating central regulatory roles, such as H4K5acK16ac or H3K9meK14ac, were more prevalent in the treated group as compared to the control (**Fig. 6E, top-right).** Taken together, the analysis of how hPTMs indirectly interact or co-regulate, and how this changes across different biological contexts, will serve as a valuable tool in deciphering the “histone code” that was hypothesized over 20 years ago^2^.

## Discussion

This study presents a novel MS-based workflow that enables comprehensive and accurate profiling of hPTMs at single-cell resolution (**Fig. 1A)**. We successfully identified 68 histone peptidoforms encompassing 25 individual hPTMs of which at least 53 were confidently quantified to enable downstream analyses. Comparatively, in bulk histone analysis via LC-MS/MS, approximately 200-300 histone peptidoforms are typically identified^28,44,46,47^. Our workflow constitutes a significant advance over previous antibody-based single-cell epigenomic approaches, which are limited in the number of targets and prone to cross-reactivity. Moreover, our methodology enables the identification and quantification of less common PTMs beyond currently available antibodies (i.e. H3K23 succinylation) or previously unrecognized modifications, (i.e. histidine methylation on H2A^48^).

A key strength of our method is the ability to detect co-occurring histone modifications, or "combinatorial histone marks", on the same peptide. Such combinatorial hPTM patterns are thought to play a crucial role in establishing distinct chromatin states and recruiting chromatin-modifying enzymes. However, accurate quantification of combinatorial hPTMs from bulk samples is challenged by cell-to-cell heterogeneity. Our schPTM method has demonstrated that histone codes, such as H3K9me2K14ac and H3K9me3K14ac, are relatively abundant in single cells, which has the potential to shed light on the role of chromatin domains decorated by both silencing and activating marks. We identified H3K9me3 as having the largest variation amongst HeLa cells in culture, which is likely due to its known role in chromatin condensation during mitosis as cells in these experiments were not synchronized by cell-cycle. In contrast, the lack of variation in H2AJK5acK9ac is poorly studied, but may represent a stable epigenetic modification in proliferating cells in culture (**Fig. 3F**). Such insights into the cell-to-cell variability of the epigenetic landscape have important implications for understanding mechanisms of cellular proliferation, differentiation, adaptation, disease progression, and aging.

Because of the single-cell resolution of our method, this allowed us to further dissect the heterogeneity of chromatin dynamics in response to the HDAC inhibitor NaBut. Although this treatment is known to induce histone acetylation to activate gene expression, we identified an unexpected subpopulation of cells which had a hPTM profile suggestive of chromatin silencing (**Figs. 5D-E**). In addition, we analyzed differential hPTM cross talk, which revealed a negative relationship between H4K5acK16ac and H3K4me that was only present in the group treated with NaBut (**Fig. 6B)**. This is suggestive of an inhibitory effect of histone H4 hyperacetylation histone H3 methylation, where H4 hyperacetylation alters the local nucleosome structure such that there is reduced accessibility of methylating enzymes targeting H3. Differential hPTM cross talk analysis has the potential to shed light on how epigenetic networks are re-wired in development and disease.

There are a few areas for improvement in the current methodology. In contrast to traditional bulk analysis, we bypassed histone extraction and prepared the contents of the whole cell to minimize adsorptive losses. Intuitively, histone isolation may lead to increased sensitivity due to lower background. However, introducing this step is infeasible for single-cell sample amounts due to the centrifugation step required^15^. In addition, bypassing histone extraction can allow for the analysis of other proteins within the same cell, providing an additional layer of information on proteomic abundances and insights into broader cellular processes and their relation to the histone code. Advances in instrumentation have recently allowed the detection of protein PTMs in using standard SCP workflows, including histone modifications due to their relatively high abundance in the cell^49,50^. While these publications are excellent proofs of concept for selected PTMs, they have intrinsic limitations for hPTM analysis, which are the reduced coverage of detectable peptides due to trypsin digestion without derivatization. Furthermore, we aim to increase the analysis throughput by optimizing the chromatographic separation and employ targeted MS acquisition, such as parallel reaction monitoring (PRM)^34,51^. We expect that very short gradients, e.g. 5 min, are feasible while maintaining similar sensitivity and potentially analyze hundreds of cells per day.^34,51^

While the intricate combinatorial complexity of the histone code allows for billions of combinations of hPTMs, the comprehensive and reproducible fragmentation patterns generated with DIA enables the relative comparison of these hPTMs across large sample numbers. However, manual peak picking (e.g. due to identical mass shifts of certain PTMs that result in isobaric peptidoforms) is a major bottleneck as it is time consuming and requires in-depth knowledge of histone peptides. At present, only a limited number of tools, such as EpiProfile^41^ which is specific to data produced by a Thermo Fisher MS instrument, can automatically process histone DIA data. Thus, the development of dedicated computational tools tailored for the analysis of histone diaPASEF data from Bruker instruments will be an important step towards increasing the accessibility and scale of this approach.

In conclusion, this study presents a robust and versatile MS-based workflow to profile the histone code at single-cell resolution. We provide unprecedented insights into the epigenetic heterogeneity within cell populations, and we envision that this method will open new avenues for investigating the role of the histone code in diverse biological processes and diseases.

## Extended Methods

### Cell culture

Low passage HeLa cells (ATCC CCL-2), were grown as adherent cultures in Eagle’s Minimum Essential Medium (ATCC 30-2003) supplemented with 10% fetal bovine serum (Thermo A5256801) and 1% penicillin-streptomycin (Thermo Fisher Scientific 15140122) in 10 cm dishes (Thermo Fisher Scientific 172931) at 5% CO_2_ and 37°C. Cells were passaged at 70-80% confluence every 3 days. For drug treatment studies, fresh media was supplemented with either 1X PBS (Thermo Fisher Scientific 10010023) as a vehicle control or sodium butyrate (Sigma-Aldrich 303410) at a concentration of 5 mM. The treatment was administered for 24 hours, after which the cells were harvested for single-cell isolation and downstream sample preparation.

### Cell isolation

Prior to harvesting, cells were washed with ice-cold 1X PBS. Cell detachment was achieved by treating the cells with 1 mL of trypLE (Thermo Fisher Scientific 12604013), followed by incubation at 37°C for 3-5 min. The cells were washed with ice-cold 1X PBS, centrifuged at 500 g for 5 min at 4°C, the cell pellet was resuspended in ice-cold 1X PBS and strained using a 40 µm cell strainer (Millipore Sigma CLS431750), followed by centrifugation and washing. Cell concentration was determined using a Countess™ 3 Automated Cell Counter (Fisher Scientific AMQAX2000) to obtain a 200 cells/µL suspension. Cells were kept on ice prior to cell sorting.

### Single-cell isolation and histone peptide preparation

All reagents were prepared fresh on the day of sample processing. A type 1 piezo dispense capillary (PDC; Cellenion) was used for cell sorting and liquid handling within the cellenONE (Cellenion). All workflow steps outside of the cellenONE were done in a sterile cell culture hood with laminar air flow. Custom scripts were written within the cellenONE software for each step of the process (see ‘Code Availability’). 1 µL of lysis reagent (0.2% dodecyl maltoside Thermo 89902 in 1M TEAB Millipore Sigma 18597) was dispensed to respective wells of 384-well plates (Thermo Fisher Scientific AB1384) using an electronic multichannel pipette (Sartorius LH-747321) and then placed in a 384-well plate holder in the cellenONE at 4°C. A 1 pg/nL bovine histone standard (Sigma-Aldrich H9250) suspension in 1X PBS was degassed on ice for 10 min, and indicated amounts (i.e. 10, 20, 40, 80, and 100 pg) were dispensed using the ‘Reagent_Dispense’ program (**Supplementary Table 2**). Following this, the plate was sealed (Thermo AB0626), vortexed for 5 sec, and centrifuged at 500 g for 30 sec. To filter out dead cells during cell sorting, DAPI stain (Thermo D1306) was added to the cell suspension prior to sorting at a concentration of 1 µg/mL, followed by degassing on ice for 10 min. Live single-cells were sorted into the plate using the ‘cellenONE_Basic_WithscanFields’ program where images of each isolated cell were acquired to verify successful isolation (**Supplementary Figs. 1A-D**). For the 100-cell sample, the original cell suspension was resuspended in lysis reagent at a concentration of 100 cell/µL and manually dispensing 1 µL into the specified wells (**Supplementary Table 2**) and were prepared identically to single-cell samples. The plate was then sealed, vortexed for 5 sec, and centrifuged at 500 g for 30 sec.

10X propionylation reagent (25% propionic anhydride Sigma-Aldrich 240311 and 75% acetonitrile Sigma-Aldrich 34851) was degassed on ice for 10 min. Nozzle parameters, i.e. voltage, pulse, and delay (400 ms), were tuned to account for low viscosity of the propionylation reagent to obtain a stable drop (**Supplementary Fig. 1E**). The ‘Derivatization_Dispense’ program was then used to dispense 100 nL of propionylation reagent to obtain a final concentration of 1X in each well. The plate was then sealed, vortexed for 5 sec, and centrifuged at 500 g for 30 sec. Images of wells were acquired to confirm successful reagent dispense (**Supplementary Figs. 1F-G**). The plate was then incubated at RT for 30 min, followed by drying inside of the cellenONE for 10-15 min at 50°C. 1 µL of master mix reagent (100 ng/µL trypsin Gold Promega V5280, 10 U/µL Benzonase® Nuclease Millipore E1014, 1% ProteaseMAX™ Promega V2071, 1 M TEAB) was dispensed into wells using an electronic multichannel pipette. The plate was then sealed, vortexed for 5 sec, and centrifuged at 500 g for 30 sec. The sealed plate was placed back into the cellenONE and incubated for 2 hrs at 50°C using the the ‘Digestion_Incubation’ program. Following the incubation, the plate was centrifuged at 500 g for 30 sec to pull down any condensation. The plate was then placed back into the cellenONE for the 2nd round of derivatization, as described above. Finally, derivatized and digested histone peptides were diluted by dispensing 2.9 µL of dilution solution (0.1% Trifluoroacetic acid Sigma-Alrich T6508, 5% DMSO Thermo Fisher Scientific 85190, mass spec grade H_2_O Sigma-Alrich 900682) using an electronic multichannel pipette. The plate was then sealed, vortexed for 5 sec, and centrifuged at 500 g for 30 sec. Prepared plates were stored at -80°C until LC-MS/MS analysis.

### LC-MS/MS analysis

Samples were acquired using a Vanquish Neo UHPLC system (Thermo Fisher Scientific) coupled to a timsTOF Ultra (Bruker Daltonik Gmbh). Peptides were separated on an Aurora Ultimate (25 cm x 75 µM, 1.7 µm particle size and 120 Å pore size IonOpticks AUR3-25075C18-CSI) integrated emitter column and eluted over a 30-minute window using the following gradient: 5-23% Solvent B (0.1% formic acid Sigma-Aldrich F0507, acetonitrile) in 25.7 minutes, 23-35% Solvent B in 2.6 minutes and 35-45% Solvent B in 1.7 minutes, at a flow rate of 200 nL/min. For shorter chromatographic gradients, peptides eluted either over a 10-minute window ranging from 5-23% Solvent B in 5.8 minutes and 23-45% Solvent B in 4 minutes or over a 5-minute window ranging from 5-45% Solvent B in 4.8 minutes. The actively acquired sample plate was stored in the Vanquish Neo autosampler at 7°C. To minimize evaporation, samples were unsealed for 12 hours of measurement time and the entire sample volume was injected for every analytical run.

For dda-PASEF experiments, full MS data were acquired in the range of 100-1700 m/z and 1.6 1/K0 [V-s/cm^-2^] to 0.6 1/K0 [V-s/cm^-2^]. Precursors were isolated with 2 m/z at 700 m/z or 3 m/z at 800 m/z. For dia-PASEF, full MS data were acquired in the range of 100-1700 m/z and 1.3 1/K0 [V-s/cm^-2^] to 0.6 1/K0 [V-s/cm^-2^]. DIA windows ranged from 300 m/z to 1000 m/z with 30 Th isolation windows and were acquired with ramp times of 200 ms. High sensitivity detection for low sample amounts was enabled without dia-PASEF data denoising. For both dia- and ddaPASEF the collision energy was ramped as a function of increasing mobility starting from 20 eV at 1/K0 [V-s/cm^-2^] to 59 eV at 1.6 1/K0 [V-s/cm^-2^].

### Histone peptide identification and quantification

Peptide identification and quantification was performed using Skyline v22.2 as previously described^12^. Briefly, ArgC was used as the digestion enzyme due to peptide derivatization. Charge states 1+ and 2+ were considered for precursor ions, and charge states 1+ and 2+ were considered for product ions. All sidechain modifications were set as variable modifications and a maximum of four modifications were allowed to occur on any peptide. Propionyl on K and the N-terminus were set as fixed. Importantly, Skyline only allows one modification per amino acid. However, a first position K can present a propionyl on the N-terminus and a variable modification on the side chain. To allow for multiple modifications per amino acid, new modifications were defined by combining the mass of propionyl groups with all possible side chain modifications. Next, a FASTA database containing all histone variants and all possible modified histone ArgC peptides were used to create a target library in Skyline, as previously described^13^. Shared peptides between different histone variants were allocated to the first occurring protein. The staggered windows used for diaPASEF acquisition as described above were set as the isolation scheme. Raw diaPASEF data of the 100-cell and 100 pg histone standard samples were used to manually curate a reference for single-cell samples. This was done based on the alignment of MS1 and MS2 XICs, relative retention time differences expected between unmodified and modified peptides, and filtering for unique transitions (if applicable). Peptides that could not be confidently identified were filtered out for further analysis. One Skyline project was used to create a spectral library to facilitate peak picking for all single-cell samples and additional separate projects per batch. Peak picking was limited to scans within 2 minutes of the MS/MS ID in the spectral library with manual adjustment if needed. Each identified peptide was validated against signal absence in ‘empty wells’ and peak alignment with the histone standard and the 100-cell samples. All curated skyline projects are accessible through the Panorama database, including spectral libraries (see ‘Data Availability’).

### Isobaric peptide deconvolution

Histone H4 contains isobaric peptides that did not separate chromatographically (**Supplementary Fig. 3A**). Thus, we deconvoluted the MS1 signal intensity using a combination of unique fragment ions for each peptidoform (**Supplementary Fig. 3B**) as has been previously described^41^. All fragment ion intensities unique for the isobaric peptide were summed and the contributing proportion of that isobaric peptide to all fragments was calculated. The relative proportion of isobaric peptides that contain a unique combination of fragment ions is calculated through their proportion relative to the sum of all the fragment ions of that peptide. The relative contribution proportion of all isobaric peptides totals to the MS1 intensity of the respective isobaric peptide. These calculations can be expressed as the following set of equations:

For a set of n isobaric peptides (P1, P2, …, Pn), where P1 has unique fragments and P2 to Pn have a unique combination of fragments:

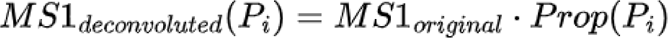

Where:

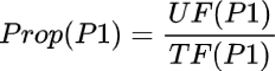

Thus, for *i* = 2 to *n*:

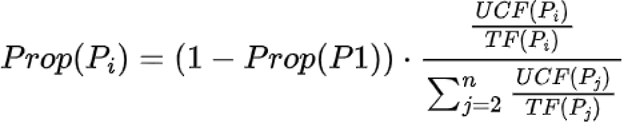

Given:

*MS1_original_* = Original (convoluted) MS1 signal

*MS1_deconvoluted_(P_i_)* = Deconvoluted MS1 signal for peptide *i*

*Prop(P_i_)* = Proportion of signal attributed to peptide *i*

*UF(P_i_)* = Sum of unique fragment intensities for peptide *i*

*UCF(P_i_)* = Sum of unique combination fragment intensities for peptide *i*

*TF(P_i_)* = Sum of all fragment intensities for peptide *i*

### Derivatization efficiency analysis

To estimate the derivatization efficiency single-cell and 10 pg bovine histone standard diaPASEF data was evaluated with the unmodified H3.1 9-17 peptide in its fully propionylated (ArgC-like peptide with propionyl modifications at K and N-terminus), the non-propionylated (Tryptic peptide with 1 or 2 missed cleavages), under-propionylated in the 1st round of derivatization (Tryptic peptide with 1 missed cleavage and propionyl modifications at K and N-terminus), under-propionylated in the 2nd round of derivatization (ArgC-like peptide with propionyl modifications at K or over-propionylated (ArgC-like peptide with propionyl modifications at T forms using Skyline^52^. The derivatization efficiency of peptides in ddaPASEF was evaluated using the software package SpectrumMill (SM), version 8.02. (Broad Institute, proteomics.broadinstitute.org). Briefly, only MS/MS spectra with precursor sequence MH+ in the range 300 to 6000 Da with a precursor charge 1 to 5 were extracted and queried using the following parameter: instrument: ESIQEXACTIVEHCD-HLA-v3; fixed modifications: propionyl-D0 on peptide N-term; variable modifications: propionylation of K with tryptic cleavage specificity allowing for a neighboring P. Matching tolerance was set to ±15 ppm for precursor and product masses and a minimum matched peak intensity of 40% against the human Uniprot database with 264 common contaminants resulting in 65068 entries. Peptide spectrum matches with <1% false discovery rate using the target decoy estimation were filtered for a minimum sequence length of 6 amino acids. PSMs were consolidated to peptides using the SM protein/peptide summary module with case-sensitive peptide-distinct mode. The percent propionylation reflects the number of propionylated peptides versus all peptides and full propionylation on internal K.

### Peptide filtering

Peptides that were only detected in their unmodified form were removed. For peptides with multiple precursor charges (i.e. +2 and +3 for H3K4un and H3K4me) the charge state with the lower median MS1 signal intensity across all cells was removed. Peptides which were not detected in greater than 50% of cells were removed. Peptides with less than 120% the median MS1 signal intensity in single-cell samples relative to empty samples were removed (**Extended data Fig. 2A, red stars**). Peptides with a median correlation of less than R=0.8 or insignificant correlation (p-value > 0.05) relative to the titration curve were removed (**Extended data Fig. 3B, red stars**).

### Cell filtering

Based on MS1 signal distributions of ‘empty wells’, peptides with a MS1 signal distribution lower than 1 median absolute deviation of the global median intensity were removed.

### hPTM relative abundance Normalization

For each modified peptide the MS1 intensity of that peptidoform was normalized to the summed MS1 intensity of all shared peptide sequences (e.g. for H2A1 4-11, H2A1K5unK9un and H2A1K5acK9un were used to create the peptide ratio) and Log(1 + *x*) transformed. Single PTM ratios were calculated by summing the peptide ratios for every unique modification at a respective amino acid within a peptide (e.g. H3K9ac = H3K9acK14un + H3K9acK14ac). Global PTM ratios were calculated by summing the PTM ratios for each unique modification (e.g. ac = H3K9ac + H3K14ac + H4K8ac etc.). The summarized H4 acetylation PTM ratios were calculated by summing the peptide ratios of peptides containing 1 (mono), 2 (di), or 3 (tri) acetylations (e.g. mono = H4K5acK8unK12unK16un + H4K5unK8acK12unK16un + H4K5unK8unK12acK16un + H4K5unK8unK12unK16ac).

### Batch correction

The presence of batch effects was assessed through the qualitative analysis of the first 2 principal components and their correlation to batch identities (e.g. **Extended Data Fig. 5A-C**). The removeBatchEffect function from the limma package^53^ was used to correct for the observed batch effect.

### Single-cell vs. 100-cell bulk comparisons

100-cell bulk and single-cell samples from batches 1-3 were processed as described above. The mean hPTM relative abundance was calculated across all single-cells (pseudobulk) and correlated the mean of the 100-cell bulk samples. hPTM relative abundance per cell was compared to the mean histone peptide ratio of 100-cell samples. PCA of single-cell and 100-cell samples were performed on normalized, batch-corrected, centered and scaled hPTM relative abundances using the PCAtools package^54^.

### Technical and Biological Noise Analysis

To define technical noise, commercially available bovine histone standard samples from batches 1-3 were analyzed identically to single-cell samples as described above. The CVs of normalized and batch-corrected or non-batch-corrected hPTM relative abundances were calculated for histone standard samples within each batch (intra-) or across batches (inter-). Single-cell level histone standards were compared to external datasets, including histone peptide ratio data of HeLa cell derived bulk technical replicates (Sidoli et al., 2015 A^32,43,44^, 5 replicates, 16 peptidoforms; Sidoli et al., 2015 B^32,43,44^, 4 replicates, 12 peptidoforms) and level 3 normalized data (Light(PTM)/Heavy(PTM) divided by Light(NORM)/Heavy(NORM)) of processing replicates for cell lines in the NIH Library of Integrated Network-Based Cellular Signatures consortium (Dele-Oni et al., 2021^32,43,44,55^, 3 replicates, 29 peptidoforms; https://panoramaweb.org/lincs_pccse_gcp_2020.url). For both external datasets, peptidoforms were matched to those identified in histone standard samples. CVs were calculated for the matched peptidoforms of the technical replicates in the external datasets relative to the inter-batch CV of the respective peptidoforms of the histone standards in the current dataset.

The CVs of normalized and batch corrected peptidoforms were calculated across all single cells in all batches. To remove expected technical noise, the variance of each peptidoform in the 10 pg bovine histone standard is subtracted. Additionally, the square root of the adjusted noise of each peptidoform is divided by its respective mean to obtain the adjusted biological CV, and peptidoforms with a greater technical than biological + technical variance were discarded.

### Cell classification analysis

Single-cells from batches 1-3 (22 control single-cells) and batches 4 and 5 (9 control, 16 treated, 33 mixed single-cells) were processed as described above to obtain normalized and batch-corrected hPTM relative abundances. PCA of single cells was performed using the PCAtools package^54^ where histone peptide ratios were centered and scaled. Following this, summarized H4 acetylation PTM ratios were calculated as described above and PCA was performed on these scaled and centered summarized ratios (Z-scores). The heatmap of scaled and centered summarized ratios (Z-scores) was plotted using the ComplexHeatmap package where hierarchical clustering was performed on the summarized H4 acetylation PTMs (rows) and single-cells (columns)^56^. To classify cells in the mixed group to the control or treated groups, we trained and evaluated a K-nearest-neighbors (KNN) classifier based on the scaled and centered summarized ratios (Z-scores). To evaluate the KNN classifier, cells from the control and treated groups were randomly split 50:50 into training and test groups where k = 2. Following this, all cells from the control and treated groups were used for training to predict the treatment group of mixed cells. The predicted labels (control or treated) of cells in the mixed group were then used in the downstream analyses.

### Differential abundance analysis

100-cell bulk and single-cell samples from batches 1-5 were processed as described above to obtain normalized and batch-corrected hPTM relative abundances. Predicted labels of cells originated from the mixed group in batches 4 and 5 were used to increase the number of cells in the control and treated groups. For ∼100-cell bulk samples, differential abundance of modified peptidoforms (4 treated vs 9 control) was performed using an unpaired two-sided T-test followed by Benjamini-Hochberg multiple hypothesis correction. For single-cell samples, differential abundance of modified peptidoforms (29 treated vs 43 control) was performed using an unpaired Wilcoxon rank sum test followed by Benjamini-Hochberg multiple hypothesis correction. Median log_2_ fold changes of significant peptidoforms per group (adjusted p-value < 0.05 and Log_2_ fold change > 0.5) were scaled and centered (Z-scores) using the ComplexHeatmap package where hierarchical clustering was performed on the peptidoforms (rows) and single-cells (columns)^56^. Differential abundance of single-cell global PTM relative abundances and summarized H4 acetylation PTM relative abundances were performed using an unpaired Wilcoxon rank sum test.

### Sub-clustering analysis

Following differential abundance analysis, the significant peptidoforms (adjusted p-value < 0.05) within cells of the treated group were scaled and centered (Z-scores) to cluster histone peptides (rows) and single-cells (columns). 2 subclusters were identified at the 2nd level of the hierarchical clustering tree which were labeled ‘A’ and ‘B.’ Differential abundance analysis of modified peptidoforms (6 subcluster A vs 23 subcluster B) was performed using an unpaired Wilcoxon rank sum test followed by Benjamini-Hochberg multiple hypothesis correction. Median log_2_ fold changes of significant peptidoforms (adjusted p-value < 0.1) per peptidoform for each group were scaled and centered (Z-scores) using the ComplexHeatmap package where hierarchical clustering was performed on the peptidoforms (rows) and single-cells (columns)^56^.

### PTM co-variation analysis

Single-cell samples from batches 1-5 were processed as described above to obtain normalized and batch-corrected hPTM relative abundances. For each group (29 treated and 43 control) and all modified peptidoforms, a covariance matrix was calculated from Pearson R and p-values of each PTM-PTM correlation using a two-sided T-test. The lower triangle of each covariance matrix was visualized using the corrplot package^57^ to only display significant relationships (p-value < 0.05). Comparison of treated vs control covariance matrices was performed using Mantel’s permutation test with 10,000 permutations, PTM-PTM correlations were followed by Benjamini-Hochberg multiple hypothesis correction. Log_2_(*x* + 1) fold changes comparing PTM-to-PTM correlations between treated vs control are displayed as a histogram. Scatter plots of peptidoform relative abundance for specific PTM-PTM relationships were fit using a linear model. In addition, Pearson R correlation coefficients and p-values of the correlations were calculated using a two-sided T-test.

### Cell-cell variation analysis

Single-cell samples from batches 1-5 were processed as described above to obtain normalized and batch-corrected hPTM relative abundances. For each group (29 treated and 43 control) and all modified peptidoforms, the pairwise Euclidean distance between all cells within each group was calculated and then compared using an unpaired T-test. Following the same procedure, the CV within each group was calculated and then compared using an unpaired T-test.

## Data availability

The raw mass spectrometry-based proteomics data generated in this study have been deposited in the MassIVE database under accession code MSV000094801 [https://massive.ucsd.edu/ProteoSAFe/dataset.jsp?accession=MSV000094801]. The data will be made publicly available after publication.

## Code availability

All analysis scripts to process data and re-create figures are available at: https://github.com/cutleraging/single-cell-histone-ptm

## Acknowledgements

We thank all members of the Sidoli and Vijg laboratories for helpful discussions. C.C. is a recipient of a SPARC Award from the Broad Institute of MIT & Harvard (#800444) that partially supported this work. L.C. acknowledges the Research Foundation Flanders – FWO for personal funding (1SF2622N) and awarding a mobility grant to join the Sidoli Lab (V400623N). This work has been supported in part by grants: T32AG023475 to R.C.; U19AG056278, U01HL145560, U01ES029519, P01AG017242, P01AG047200, RF1AG068908, P30AG038072, US Department of Defense (BC180689P1) to J.V. This work was also supported in part by grants P01CA206978 to S.A.C from the NIH, U24CA270823, U01CA271402 to S.A.C., as well as a grant from the Dr. Miriam and Sheldon G. Adelson Medical Research Foundation. This work was also supported by the Michael J. Fox Foundation and the Paul F. Glenn Center for the Biology of Human Aging at Albert Einstein College of Medicine. The Sidoli lab gratefully acknowledges for funding the Hevolution Foundation, the Einstein-Mount Sinai Diabetes center, Merck, Relay Therapeutics, and the NIH Office of the Director (S10OD030286).

## Author contributions

R.C. and S.S. conceptualized the study. R.C., C.C., M.P. and S.S. designed the experiments. R.C., L.C., and J.C., designed the sample preparation method and prepared the samples. C.C., S.V. and M.P. designed and optimized MS acquisition methods. C.C. acquired the samples. R.C., L.C., and C.C. analyzed the data and wrote the manuscript. D.D. acquired funding. S.A.J.V., M.D., J.V., M.P., S.A.C. and S.S. supervised the research and revised the manuscript.

---

**Extended Data Fig. 1.**
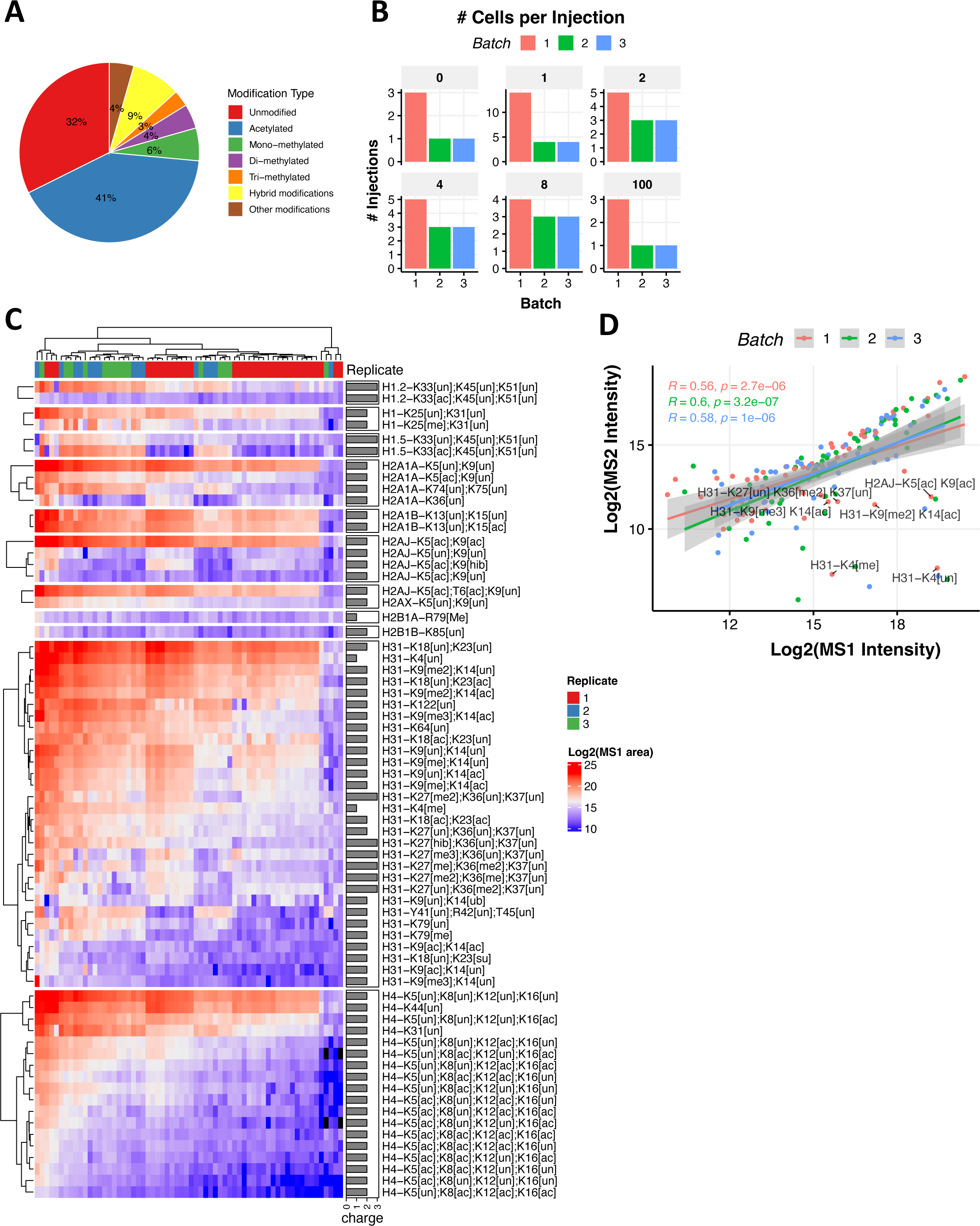
Modification types and sample information for benchmarking experiments, related to Figure 1. **A)** Proportion of unique histone PTMs calculated from peptidoforms identified from batches 1-3. The hybrid modifications are peptides that contain a combination of different types of post-translational modifications, such as methylation and acetylation. **B)** Number of injections for each cell input titration for batches 1-3. The ‘0’ sample represents the ‘empty well’ and the ‘1’ is a single cell, etc. **C)** Heatmap of MS1 intensity for identified peptidoforms in single cells. Rows ordered by histone protein and hierarchically clustered. Cell hierarchically clustered in columns. Missing values are indicated by black. **D)** Correlation analysis of MS1 and summed MS2 intensities for each peptide in each batch.

**Extended Data Fig. 2.**
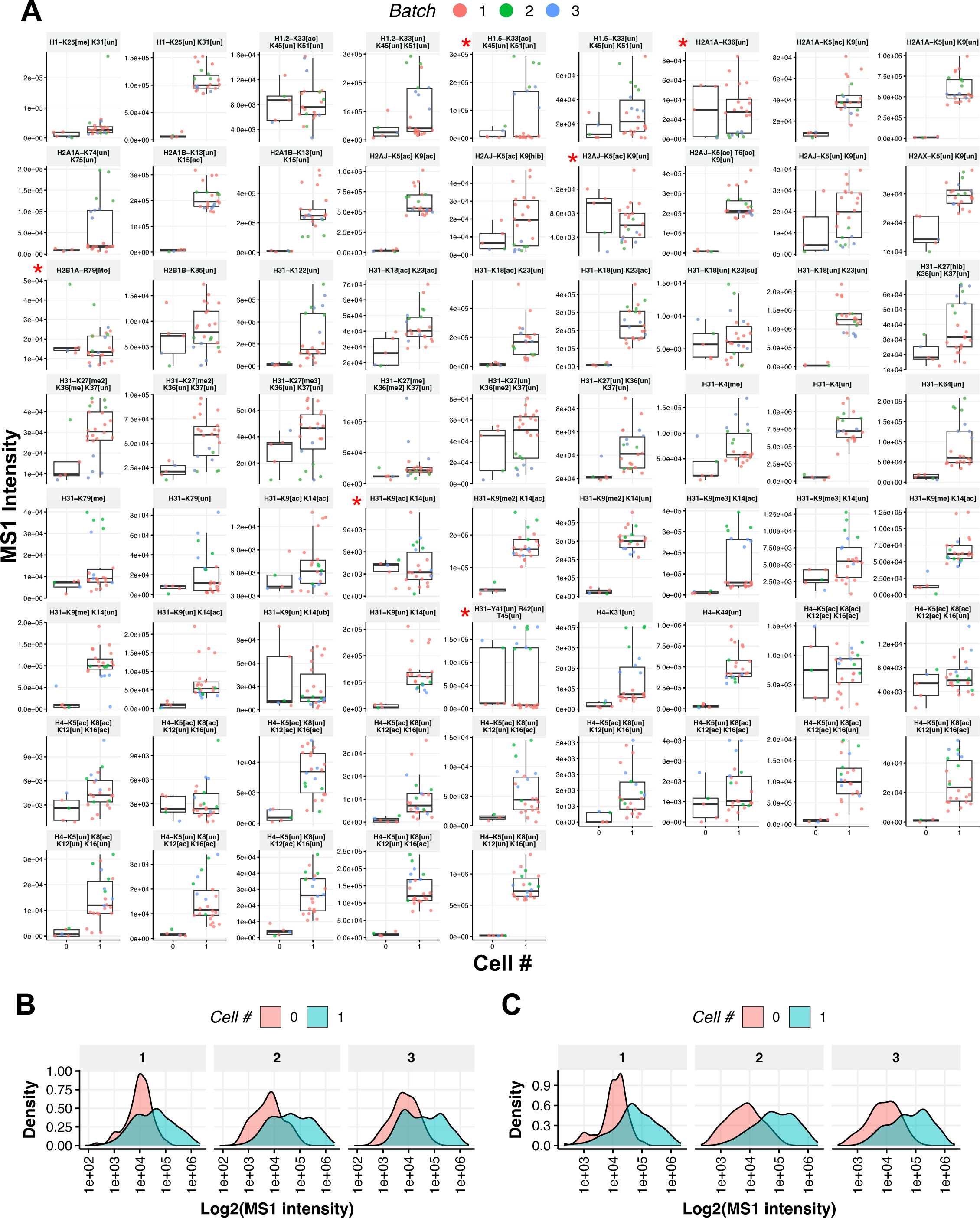
Extended analysis of background noise, related to Figure 2. **A)** MS1 intensity of all histone peptides detected in 0 cell (empty wells) and 1 cell injections from batches 1-3. The red star, when present in the top left corner of a peptide plot, indicates that the respective peptide did not pass the background noise filter and was therefore filtered out from subsequent analysis (**Methods**). **B)** MS1 intensity distributions for each batch before filtering and **C)** after removal of peptidoforms that did not pass the filtering criteria (i.e. indicated by red star in (**A**).

**Extended Data Fig. 3.**
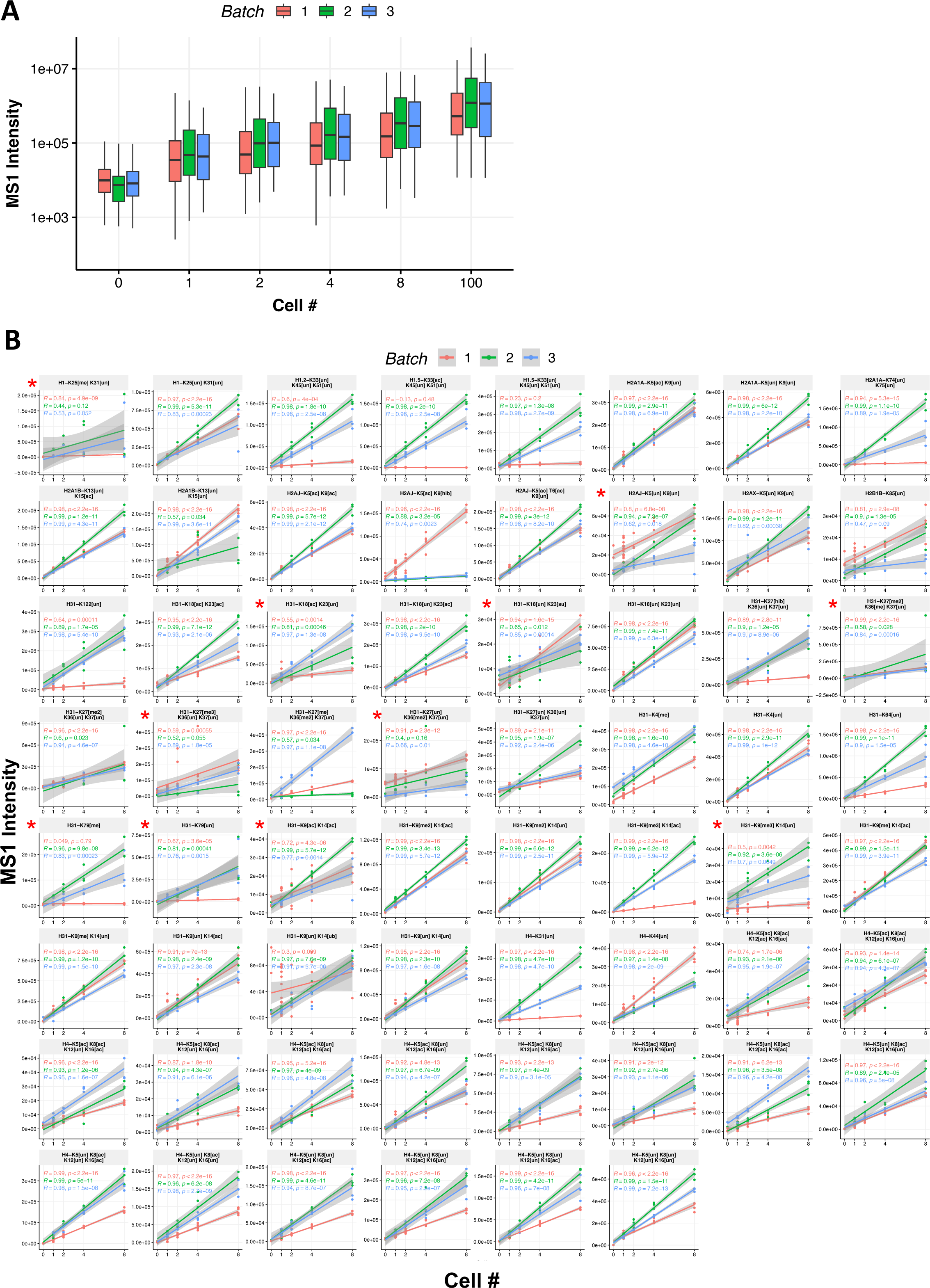
Extended analysis of cell titration curves, related to Figure 2. **A**) Global distribution of MS1 signal intensities when titrating increasing numbers of cells prepared as a single sample for batches 1-3. **B**) MS1 cell titration curves per histone peptide for batches 1-3. The red star in the top left corner of each peptide plot indicates that the respective peptide did not pass the quantitative linearity filter in relation to the control titration curves and was therefore filtered out from subsequent analysis (**Methods**).

**Extended Data Fig. 4.**
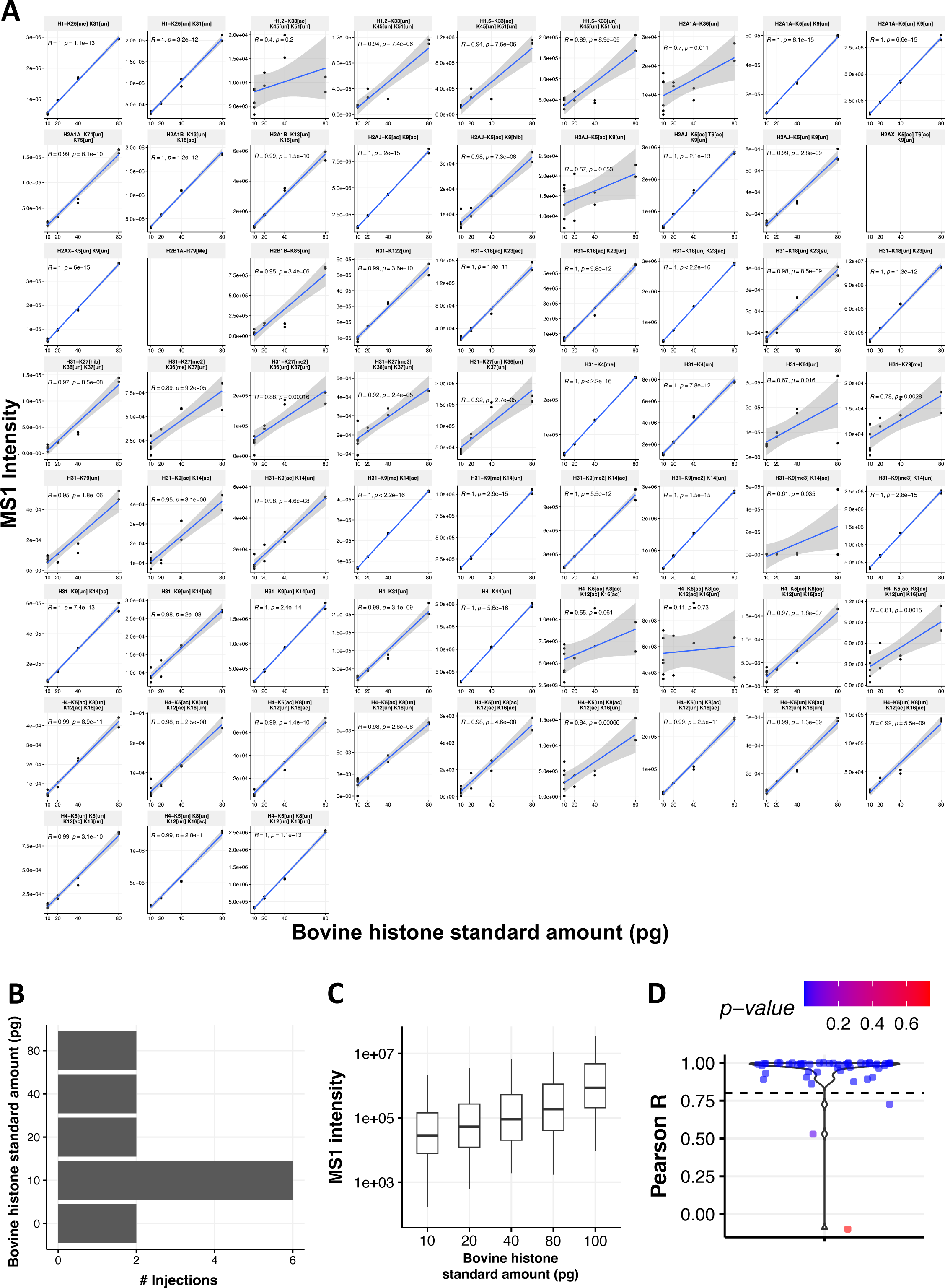
Histone standard titration curves, related to Figure 2. **A**) MS1 titration curves of bovine histone standards from replicate batch 1. **B**) Number of samples for varying amount of histone standard from replicate batch 1. **C**) Global distribution of MS1 signal intensities for samples prepared with increasing amounts of bovine histone standard from replicate batch 1. **D**) Distribution of Pearson correlation coefficients representing linearity of titration curves. Statistics: Significance of Pearson R correlation coefficients for each batch was tested using a t-test.

**Extended Data Fig. 5.**
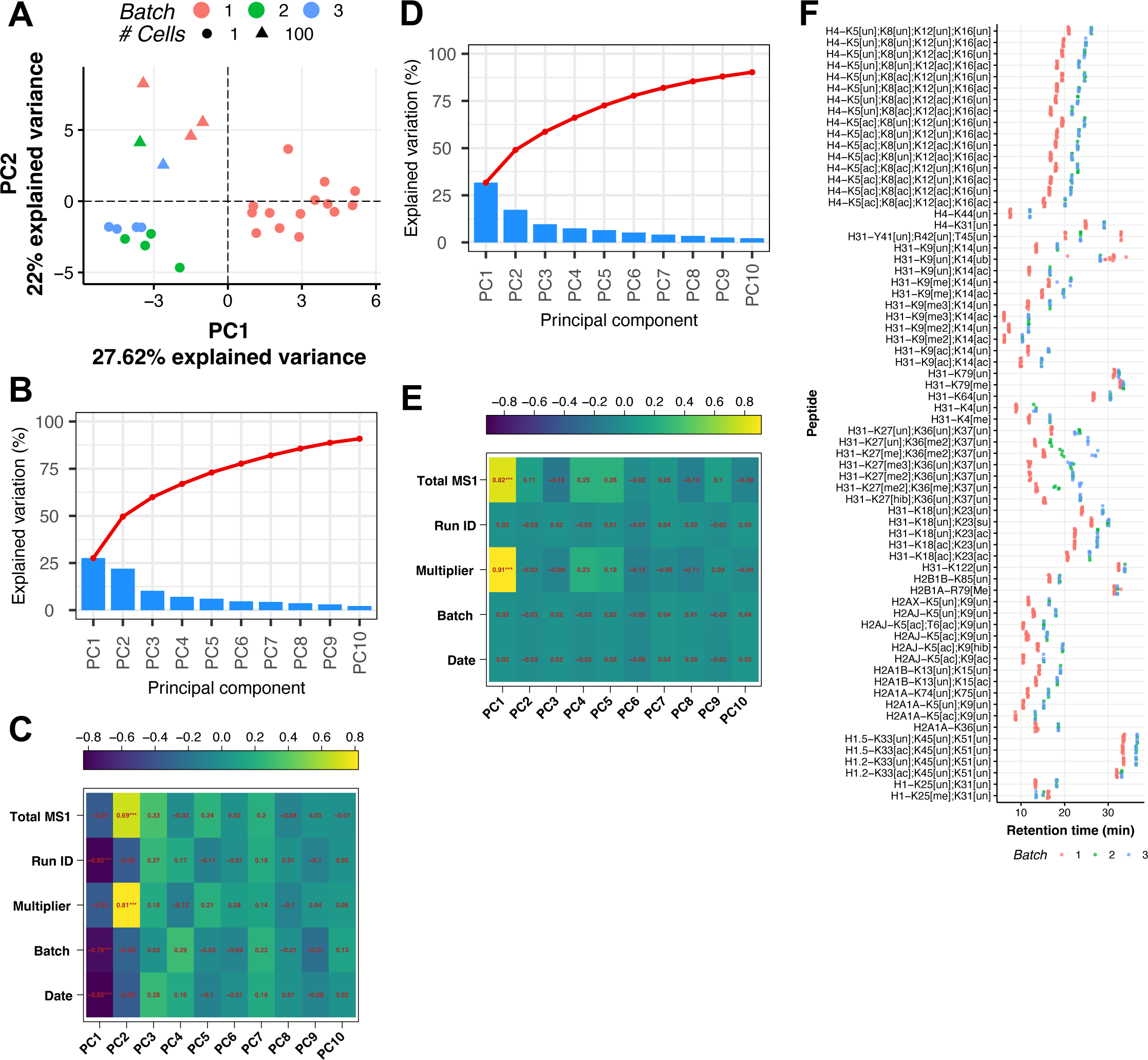
Quality control and batch-correction for samples used in benchmarking, related to Figure 2. **A)** Principal component analysis of histone peptidoform relative abundances for single-cell and 100-cell bulk samples before batch correction. **B)** Cumulative proportion of explained variation across principal components before batch correction. **C)** Correlation of each principal component to experimental variables before batch correction, showing significant correlation of PC1 with batch variable. Legend above plot represents R value. Statistics: Significance of Pearson R correlation coefficients for each batch was tested using a t-test, *** p <= 0.001. **D)** Same as (**B**) but after batch correction. **E)** Same as (**C**) but after batch correction, showing no correlation of batch variable with principal components. **F)** Retention times of filtered peptidoforms for batches 1-3. Note that batch 1 retention times are earlier due to liquid chromatography instrument calibration between batches 1 and 2-3.

**Extended Data Fig. 6.**
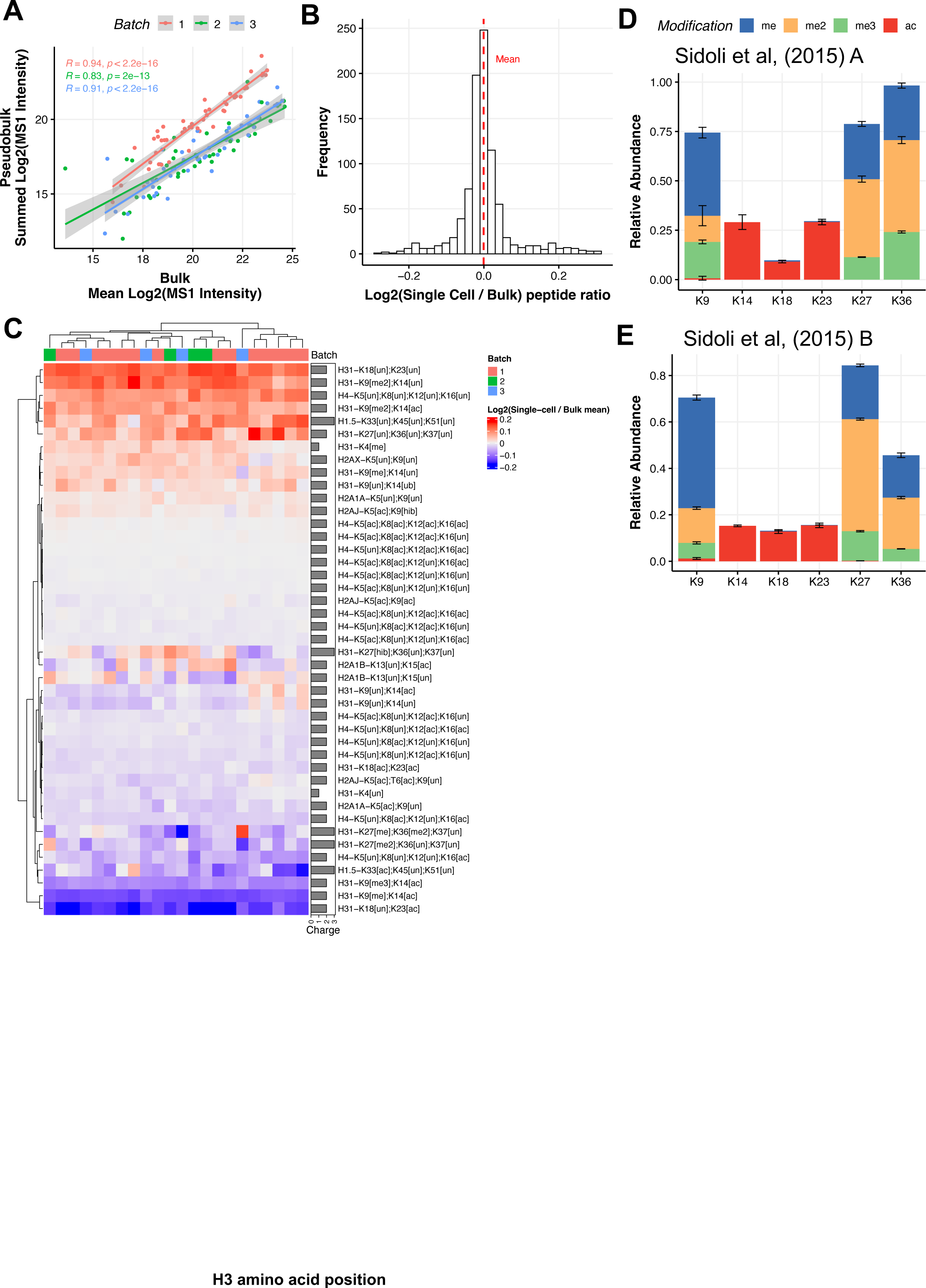
Extended comparisons of hPTM quantification in single cells with bulk samples, related to Figure 2. **A**) Correlation of histone peptidoform Log_2_ MS1 intensity between bulk (∼100-cell) and ‘pseudobulk’ (mean of single cells) samples in baches 1-3. Statistics: Significance of Pearson R correlation coefficients for each batch was tested using a t-test. **B-C**) Distribution (**B**) and heatmap (**C**) of Log_2_ fold changes resulting from comparison of histone peptidoform Log_2_ relative abundances between each single cell in batches 1-3 and average of bulk samples. **D-E**) Individual histone PTMs (single and combinatorial PTMs) relative abundances detected on histone H3 in bulk HeLa cell samples from Sidoli et al, 2015 A^43^ (**D**), and Sidoli et al, 2015 B^44^ (**E**). These are in comparison to the single-cell data shown in **Fig. 2G**.

**Extended Data Fig. 7.**
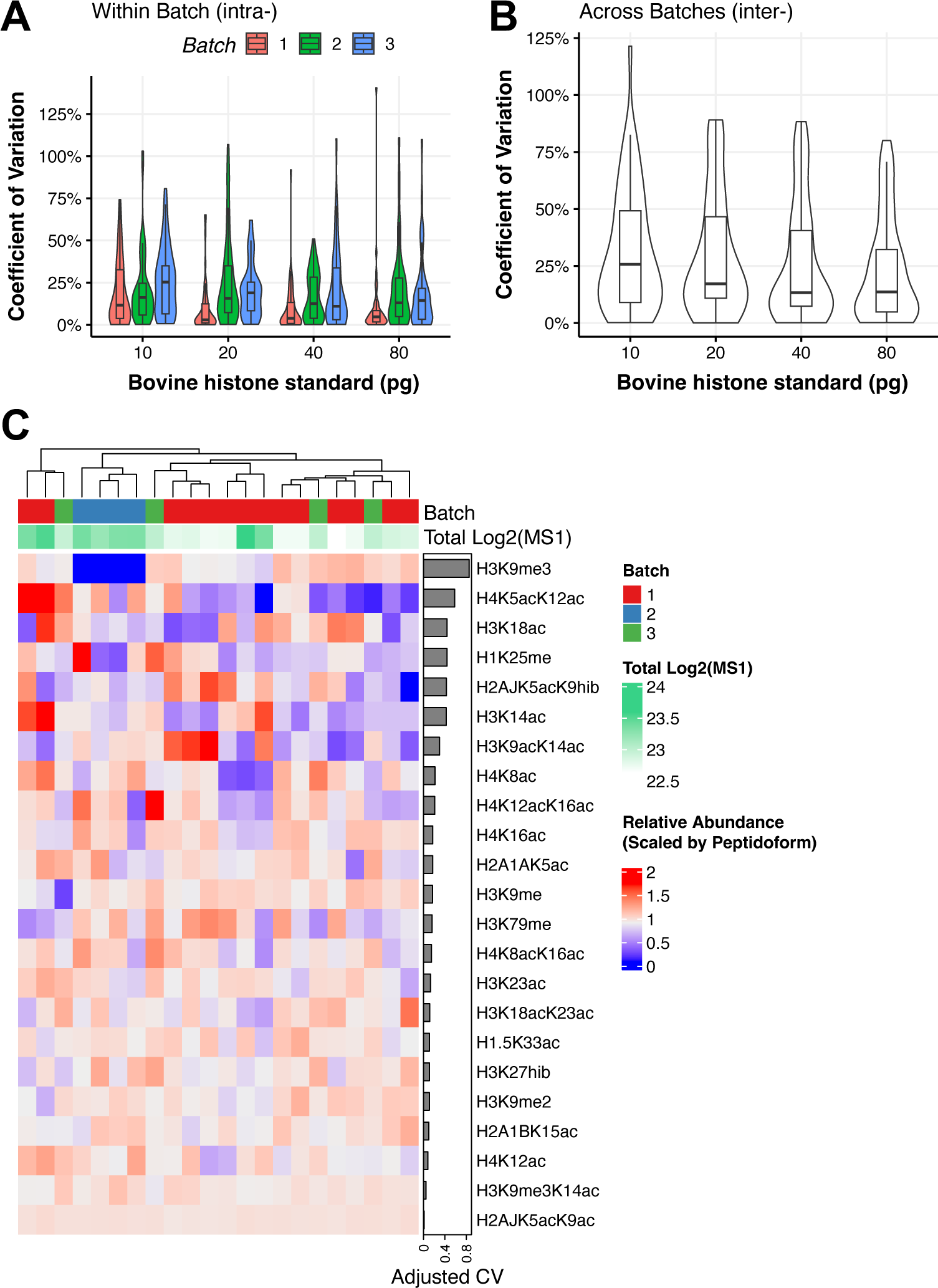
Extended analysis of technical noise in hPTM quantification, related to Figure 3. **A)** Coefficient of variation (CV) of MS1 intensities for bovine histone standards technical replicates within each batch or **B)** across batches 1-3. **C)** Heatmap of relative abundances scaled to each peptidoform organized from top-down by greatest to least CV, showing variation of peptidoform abundances across single cells.

**Extended Data Fig. 8.**
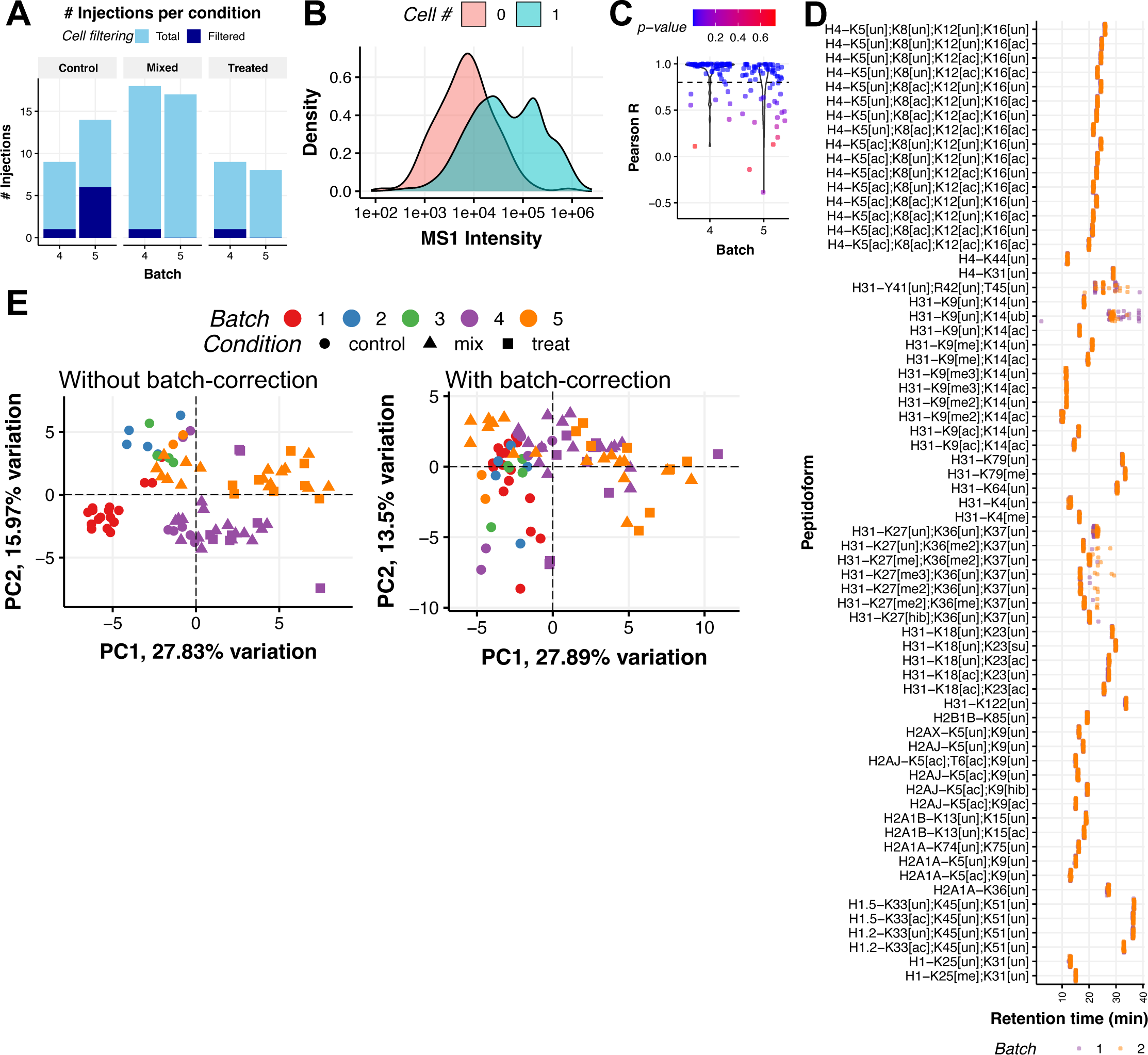
Quality control and batch correction for samples used in HDAC inhibitor treatment experiment, related to Figure 4. **A)** Number of injections (single cells) passing filtering (**Methods**) in batches 4-5 across different conditions. **B)** Distribution of raw Log_2_ MS1 intensities for all histone peptidoforms in batches 3-4 for empty wells and wells containing single cells after filtering out peptides with low signal (**Methods**). **C)** Distribution of Pearson R correlation coefficients for titration curves of all histone peptidoforms in batches 4-5. A dashed line is drawn at R=0.8 to indicate the threshold used for filtering out histone peptidoforms deemed to be non-quantifiable. Statistics: Significance of Pearson R correlation coefficients for each batch was tested using a t-test. **D)** Retention times of filtered peptidoforms for batches 1-3. **E)** Principal component analysis of histone peptidoform relative abundances for single cell samples from all batches without batch correction (left) and with batch correction (right).

**Extended Data Fig. 9.**
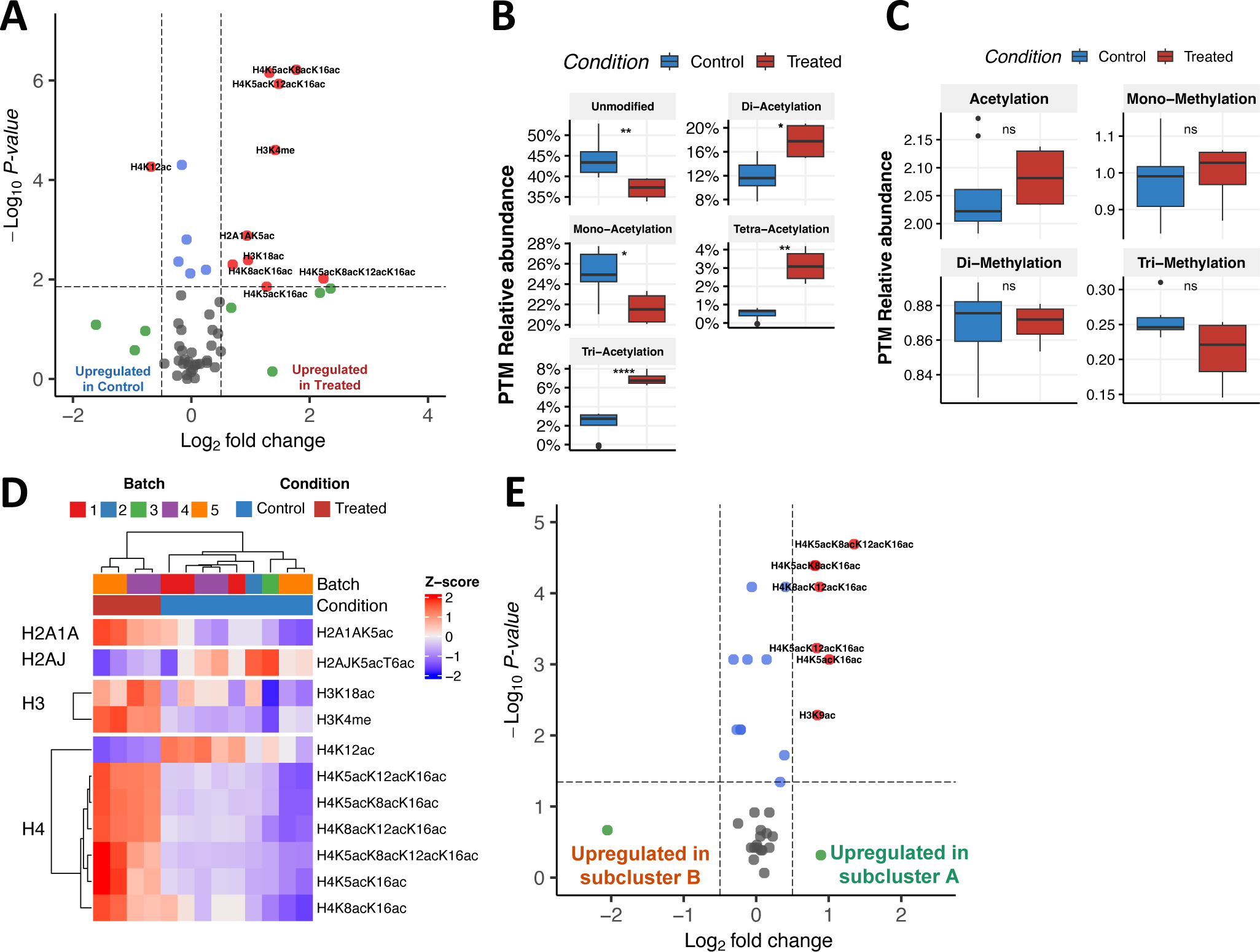
Extended analysis of HDAC inhibitor treatment, related to Figure 5. **A)** Differential abundance of modified histone peptidoforms relative abundances in 100-cell bulk samples for treated vs. control comparison. Statistics: Wilcox test with Benjamini & Hochberg multiple testing correction. Red points are peptidoforms that are significant (adjusted p-value < 0.05) and Log_2_ fold change > 0.5. Blue points are peptidoforms that are significant only. Grey points are peptidoforms that are not significant (adjusted p-value > 0.05) and Log_2_ fold change < 0.5. For table of full results, see **Supplementary Table 7**. **B)** Summarized relative abundances of global PTMs in 100-cell bulk samples for control and treated groups. Statistics: Wilcox test, ns p > 0.05. **C)** Summarized relative abundances of H4 peptide acetylation in 100-cell bulk samples for control and treated groups. Statistics: Wilcox test, * p <= 0.05, ** p <= 0.01, **** p <= 0.0001. **D)** Heatmap and hierarchal clustering of modified histone peptidoform scaled relative abundances (Z-score) that were significantly differentially abundant (adjusted p-value < 0.05) between treated vs. control groups as shown in (**A**). Cells are shown in columns and histone peptidoforms are shown in rows. Peptidoforms are grouped by histone proteins Note that no variation in histone peptidoform abundance can be seen in the treated group (right side). **E)** Differential abundance of modified histone peptidoforms relative abundances in single cells for subcluster A vs. subcluster B comparison, related to **Fig. 5E**. Statistics: Wilcox test with Benjamini & Hochberg multiple testing correction. Red points are peptidoforms that are significant (adjusted p-value < 0.05) and Log_2_ fold change > 0.5. Blue points are peptidoforms that are significant only. Grey points are peptidoforms that are not significant (adjusted p-value > 0.05) and Log_2_ fold change < 0.5. For table of full results, see **Supplementary Table 8**.

**Extended Data Fig. 10.**
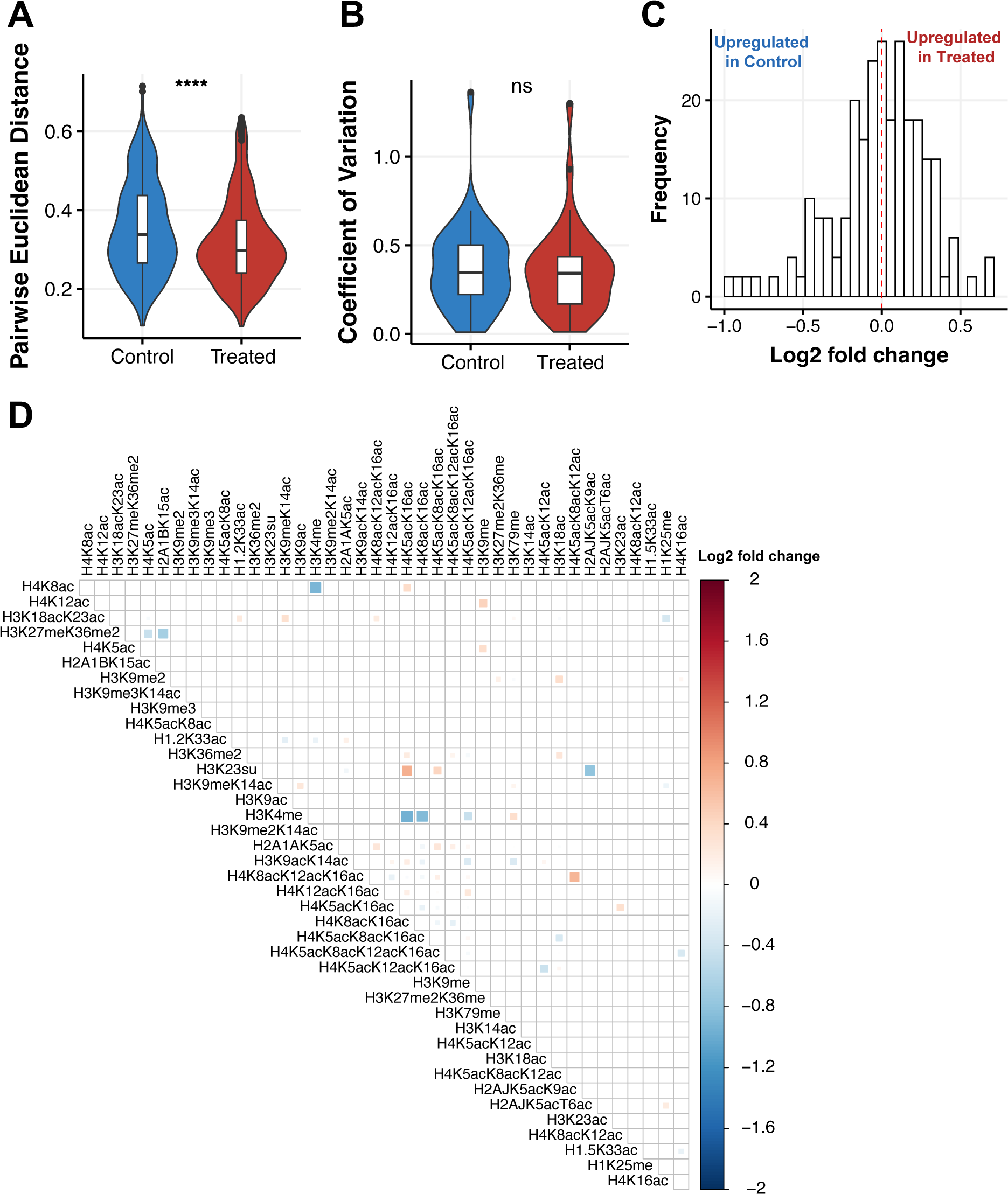
Extended analysis of histone PTM-PTM coregulation, related to Figure 6. **A)** Assessment of cell-to-cell variation by pairwise Euclidian distance (**Methods**) for control and treated groups. Statistics: Wilcox test, **** p <= 0.0001. **B)** Assessment of histone PTM noise by comparing distributions of coefficient of variation for peptidoform relative abundances between control and treated groups. Statistics: Wilcox test. **C)** Distribution of Log_2_ fold changes resulting from differential PTM-PTM correlations between treated and control groups. Pearson R correlation coefficients derived from modified histone PTM-PTM correlations were used for the comparison. Note the positive shift in the distribution, indicating stronger PTM-PTM correlations in the treated group as compared to the control group. **D)** Detailed results of differential PTM-PTM correlations as shown in (**C**). Statistics: Significance of differential Pearson R correlation coefficients was tested using a permutation test with 10,000 permutations (**Methods**). Blank cells indicate non-significant p-values.

