## Supplementary figures for "Mass Spectrometry-based Profiling of Single-cell Histone Post-translational Modifications to Dissect Chromatin Heterogeneity"

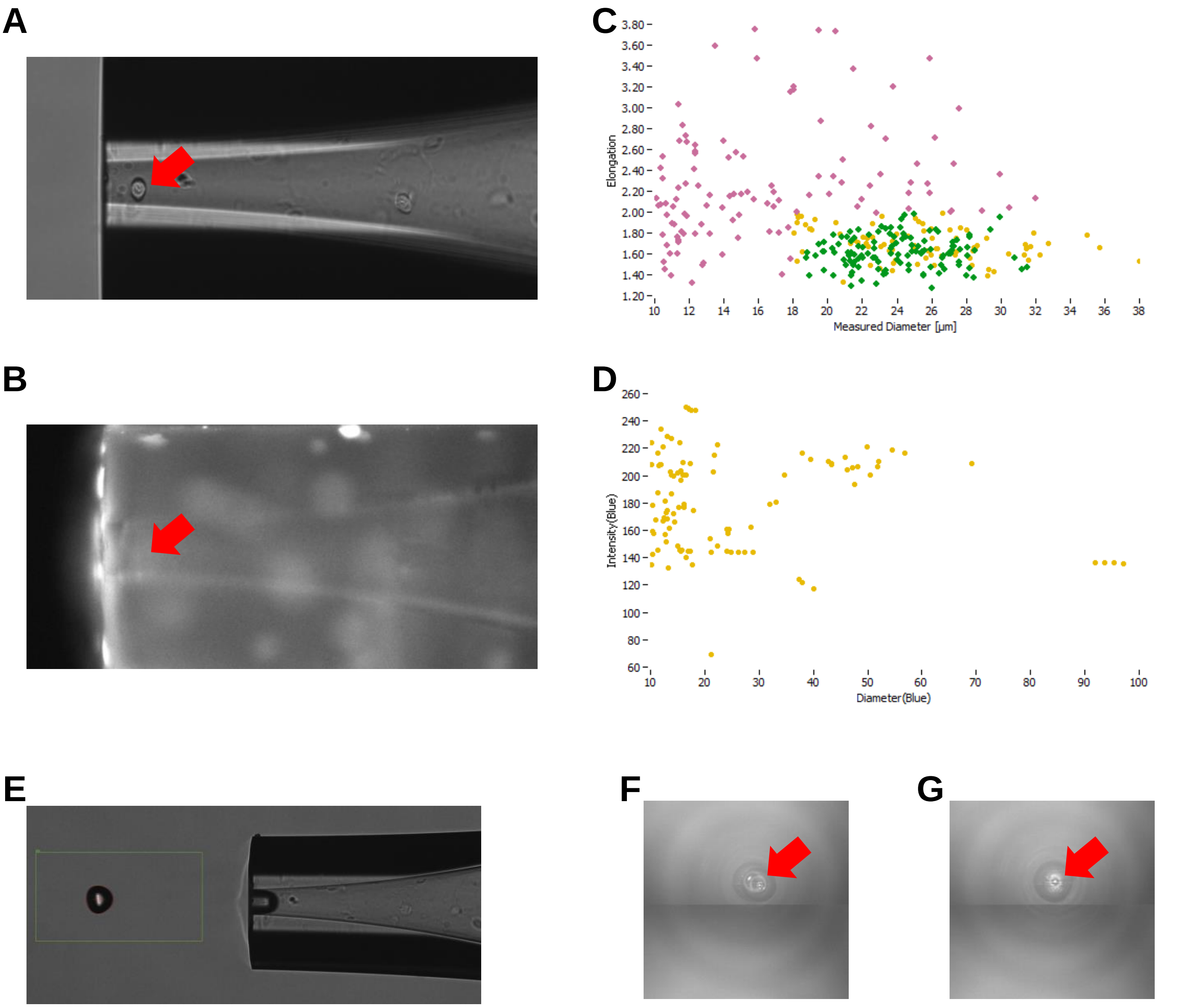


**Supplementary Fig. 1**

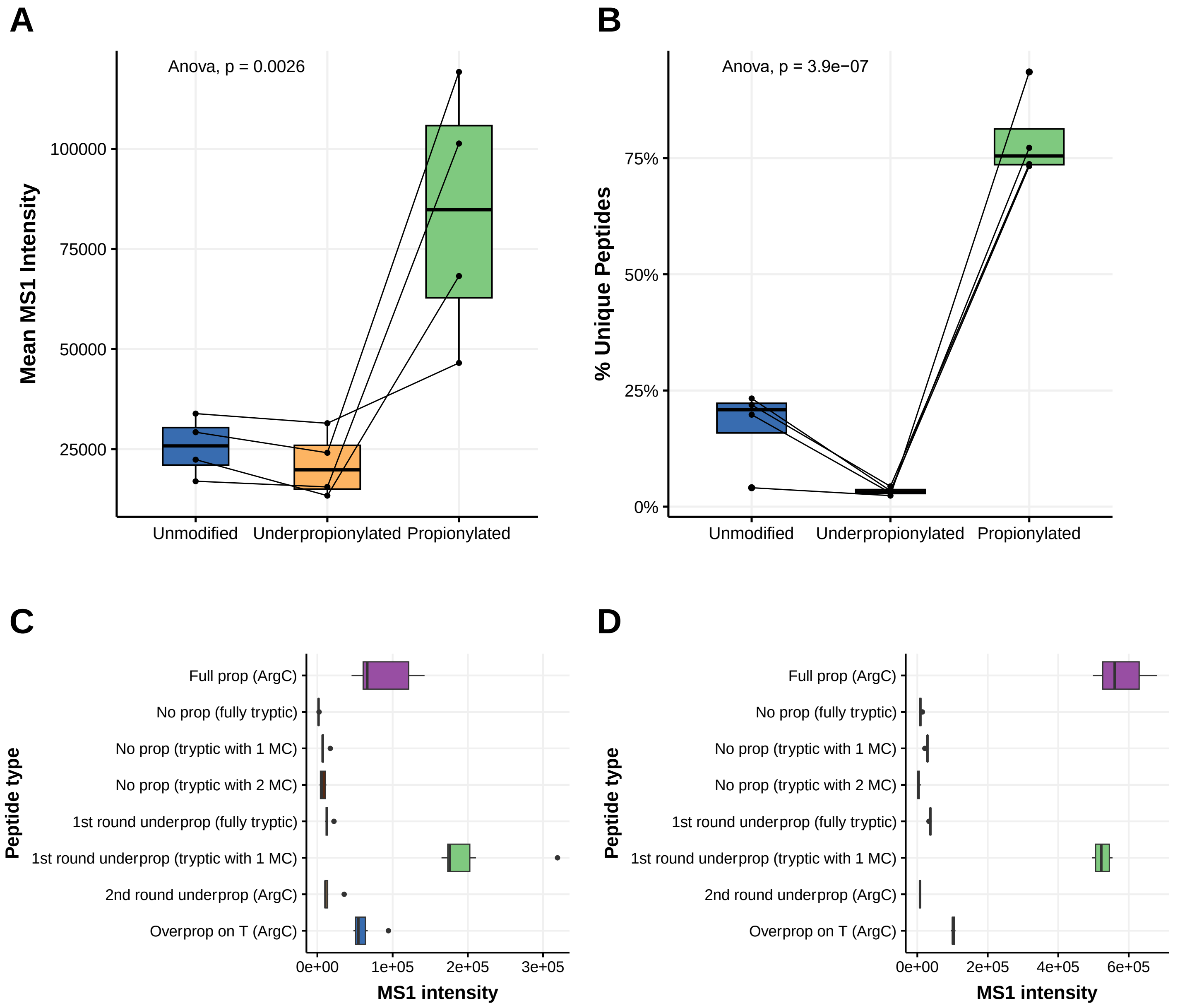


**Supplementary Fig. 2**

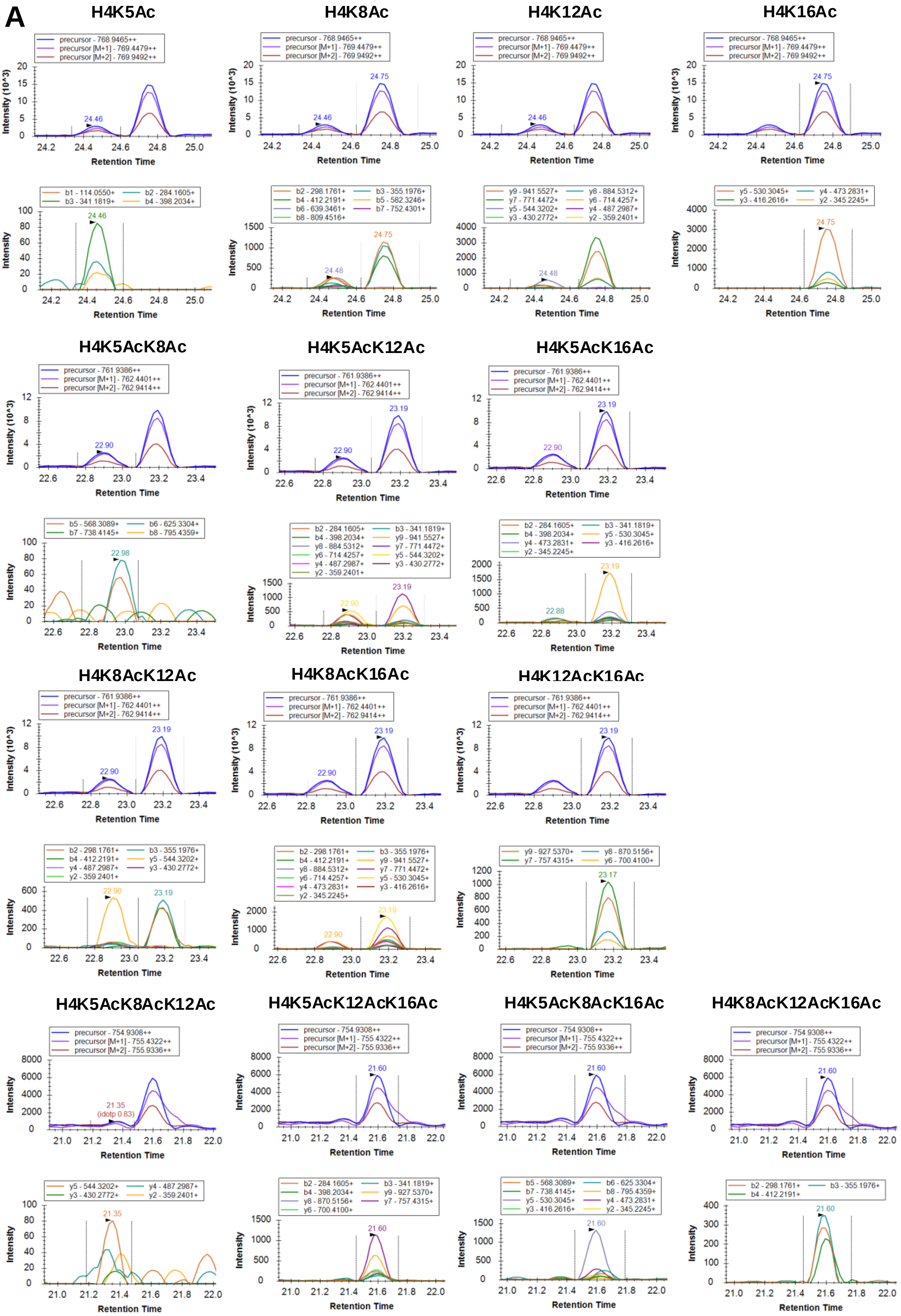


Supplementary Fig. 3

**B**

| b1 | b2 | b3 | b4 | b5 | b6 | b7 | b8 | b9 | b10 | b11 | b12 | b13 | b14 |
|---|---|---|---|---|---|---|---|---|---|---|---|---|---|
| y14 | y13 | y12 | y11 | y10 | y9 | y8 | y7 | y6 | y5 | y4 | y3 | y2 | y1 |
| G | K5 | G | G | K8 | G | L | G | K12 | G | G | A | K16 | R |

| Modification | m/z | RT | Residue |
|---|---|---|---|
| Un | 775.9543 | 26.00 | |
| Mono | 768.9465 | 24.46 | K5 |
| | | 24.48 | K8 |
| | | 24.43 | K12 |
| | | 24.75 | K16 |
| Di | 761.9386 | 22.98 | K5K8 |
| | | 22.88 | K5K12 |
| | | 22.90 | K8K12 |
| | | 23.17 | K5K16 |
| | | 23.19 | K8K16 |
| | | 23.17 | K12K16 |
| Tri | 754.9308 | 21.35 | K5K8K12 |
| | | 21.60 | K5K12K16 |
| | | 21.64 | K5K8K16 |
| | | 21.60 | K8K12K16 |
| Tetra | 747.923 | 20.00 | K5K8K12K16 |

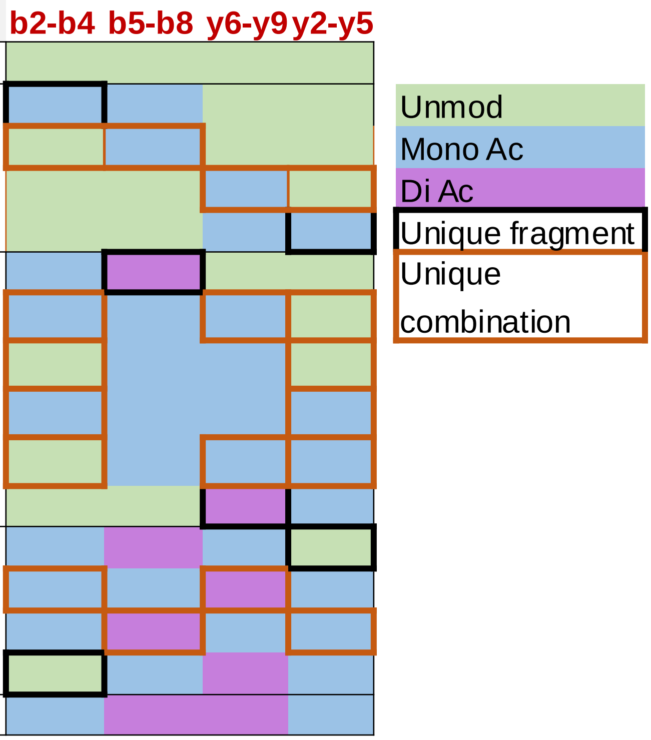


**Supplementary Fig. 3**
