## Supplementary figure legends for "Mass Spectrometry-based Profiling of Single-cell Histone Post-translational Modifications to Dissect Chromatin Heterogeneity"

**Supplementary Fig. 1 | CellenONE Cell Isolation and Reagent Dispensing Quality Control. A-B)** Brightfield (**A**) and blue fluorescence (excitation 380 nm, emission 410-450 nm) (**B**) images of a single live HeLa cell in the tip of the piezo dispenser capillary of the cellenONE as it is being isolated into the well of a 384-well plate. **C)** Plot of cells elongation and diameter of cells that were observed during cell isolation for batch 1. Green indicates cells that were isolated, pink indicates cells that did not fall into the chosen gates and were not isolated, yellow indicates cells that did fall into the chosen gates but were not isolated either due to high DAPI signal or presence of doublet. **D)** Plot from same isolation as in (**C**), but for the blue fluorescence channel showing cells that were DAPI+ and not isolated. **E)** Image of derivatization reagent (25% propionic anhydride and 75% acetonitrile) being dispensed with stable droplet. Voltage is set to 63 V, pulse is set to 50 μs, and delay is set to 400 μs. **F-G)** Brightfield images of a single-well of a 384-well plate before (**F**) and after **(G**) the dispense of 100 nl of derivatization reagent into a well containing 1 μL lysis reagent and a single-cell.

**Supplementary Fig. 2 | Propionylation efficiency in single-cells and 10 pg histone standards. A)** Mean raw MS1 intensity of different variations of peptide modification of histone peptides from 4 single-cells in batch 1 acquired with DDA (**Methods**). 20 peptides on average were identified as unmodified, 7 peptides on average were identified as under-propionylated, and 150 peptides on average were identified as fully propionylated. **B)** Unique percentage of various types of histone peptides from data shown in (**A**). **C-D)** Raw MS1 intensity of different variations of peptide modification or digestion outcomes for histone peptide H3.1 9-17 (**Methods**) from 6 single-cells (**C**) or 6 10 pg histone standard samples (**D**) in batch 1 acquired with DIA.

**Supplementary Fig. 3 | Histone H4 Isobaric Peptidoform Deconvolution**. **A)** Extracted Ion Chromatograms (XICs) showing (combination of) unique transitions for all isobaric peptidoforms on the H4(4-17) peptide. **B)** Table illustrating the unique fragment ion combinations which were used to deconvolute the MS1 signal intensity using the fragment ion intensities for each of the H4(4-17) isobaric peptides. See methods for more details.
